# Properties of Alzheimer’s disease brain-derived tau aggregates define tau processing by human astrocytes

**DOI:** 10.1101/2024.09.04.610753

**Authors:** Matthew J. Reid, Melissa Leija Salazar, Claire Troakes, Steven Lynham, Deepak P. Srivastava, Beatriz Gomez Perez-Nievas, Wendy Noble

**Author notes:** Corresponding author: Professor Wendy Noble.

## Abstract

The templated misfolding of tau proteins accounts for tau pathology spread in Alzheimer’s disease (AD). Post-translational modifications, including phosphorylation at specific residues, are closely linked with tau seeding ability and clinical disease progression. Increasing evidence supports a contributing role for astrocytes in tau spread. This study demonstrates that tau aggregates from postmortem AD brain are internalized and processed by human astrocytes. Differences in the efficiency of tau internalization, clearance and/or seeding were noted, which may reflect molecular properties of tau. Notably, we observed a direct relationship between alterations in tau handling by astrocytes and astrocyte responses, which were evident in transcriptomic data. Dysregulated genes include several previously identified as upregulated in reactive astrocytes in AD brain, as well as being implicated in pathological tau clearance by autophagy and other pathways. The study provides new insights into the complex interplay between tau molecular diversity and astrocyte responses in AD.

## Introduction

Altered interactions between neurons and glia contribute to the development and progression of many neurodegenerative diseases, including Alzheimer’s disease (AD), in which the prion-like spread of protein aggregates accompanies disease worsening. Glial cell function is altered in response to Aβ and tau aggregates (Brandebura et al., 2023; Ferrari-Souza et al., 2022), and understanding the consequences of these alterations may be key to determining the cellular mechanisms that drive the spread of pathological forms of tau through affected brain regions.

Tau forms the main component of neurofibrillary tangles in AD, as well as distinct neuropathological hallmarks in other tauopathies (Crowther & Goedert, 2000). The extensive modifications of tau in AD brain are now largely well understood (Wesseling et al., 2020), as are the structure of tau aggregates in AD brain (Fitzpatrick et al., 2017; Simic et al., 2016). However, there is considerable variation in the molecular profile of tau between individuals, with modifications such as phosphorylation at specific residues being associated with tau seeding efficiency and clinical disease progression (Dujardin et al., 2020). This reflects evidence that several tau “strains” exist in each tauopathy, with variation both within and between tauopathy groups (Sanders et al., 2014). Whether and how molecular diversity of tau affects glia in AD is not fully understood.

Astrocytes are the most abundant cell type of the brain (Miller, 2018), providing homeostatic support for neurons and helping to regulate synaptic activity (Chai et al., 2017; Santello et al., 2019) amongst other functions. In AD, heterogeneous astrocyte responses are determined by spatiotemporal stage (Serrano-Pozo et al., 2022). Astrocytes associate with neurofibrillary tangles as well as Aβ plaques (Serrano-Pozo et al., 2011), and harbor tau inclusions in the dentate gyrus (Richetin et al., 2020) which may highlight a potential role of these cells in tau spread. Indeed, astrocytes accumulate pathological tau following its spread from neurons in mice (de Calignon et al., 2012). Astrocyte phagocytic ability (Konishi et al., 2022), allows them to indirectly internalize large tau fibrils along with dead or dying neurons (Mothes et al., 2023), and while the ability of astrocytes to directly internalize synthetic tau fibrils has also been demonstrated (Martini-Stoica et al., 2018; Perea et al., 2019), their conformational structure is now understood to be different from that of AD and other tauopathies (Shi et al., 2021).

The aim of this study was to determine if the molecular properties of AD brain -derived tau influence the rate of human tau aggregate uptake by human astrocytes, and/or affect the function of astrocytes. We found tau was internalized by astrocytes, but that the rate of uptake and clearance varied for tau isolated from different AD cases. The molecular properties of tau, as defined through analysis of post-translational modifications, showed associations with both the rate of tau uptake and either seeding of endogenous tau or clearance by human astrocytes, as well as changes in astrocyte function indicated by gene expression changes. These data suggest that astrocytes also make an important contribution to the rate of tau spread in AD, which may be further confounded by changes in astrocyte function.

## Results

### 1. Tau accumulates in astrocytes in AD temporal cortex

Although the accumulation of disease-associated forms of tau in astrocytes is a characteristic feature of several primary tauopathies, it is not commonly described in AD (Kovacs, 2020; Reid et al., 2020). However, tau accumulation within hilar astrocytes of the dentate gyrus in AD brain was recently shown to mediate neuronal dysfunction and cognitive decline (Richetin et al., 2020). We examined the association of AT8+ve tau with astrocytes in Braak stage V-VI AD brain (n=6) relative to control (non-neurologically impaired, (n=3)) temporal cortex (BA21) grey matter. AD tissues showed the expected accumulation of sarkosyl-insoluble tau aggregates when examined biochemically (Fig. S1a). AT8+ tau structures were commonly detected within AD tissue, and to a lesser extent in control tissues (Fig.1a) and while most showed little colocalization with astrocytes, some AT8+ structures in AD sections were identified in glial fibrillary acidic protein (GFAP) and S100 calcium binding protein-b (S100b) -labelled astrocytes (Fig. 1a). The average AT8 levels in AD astrocytes were significantly higher than in control cases. Although there was significant variation between individual cases (F (8, 71722) = 5229, p<0.0001, R^2^=0.3684) (Fig. S1b), these data suggest that some disease-modified tau is present in astrocytes in AD brain.

**Figure 1.**
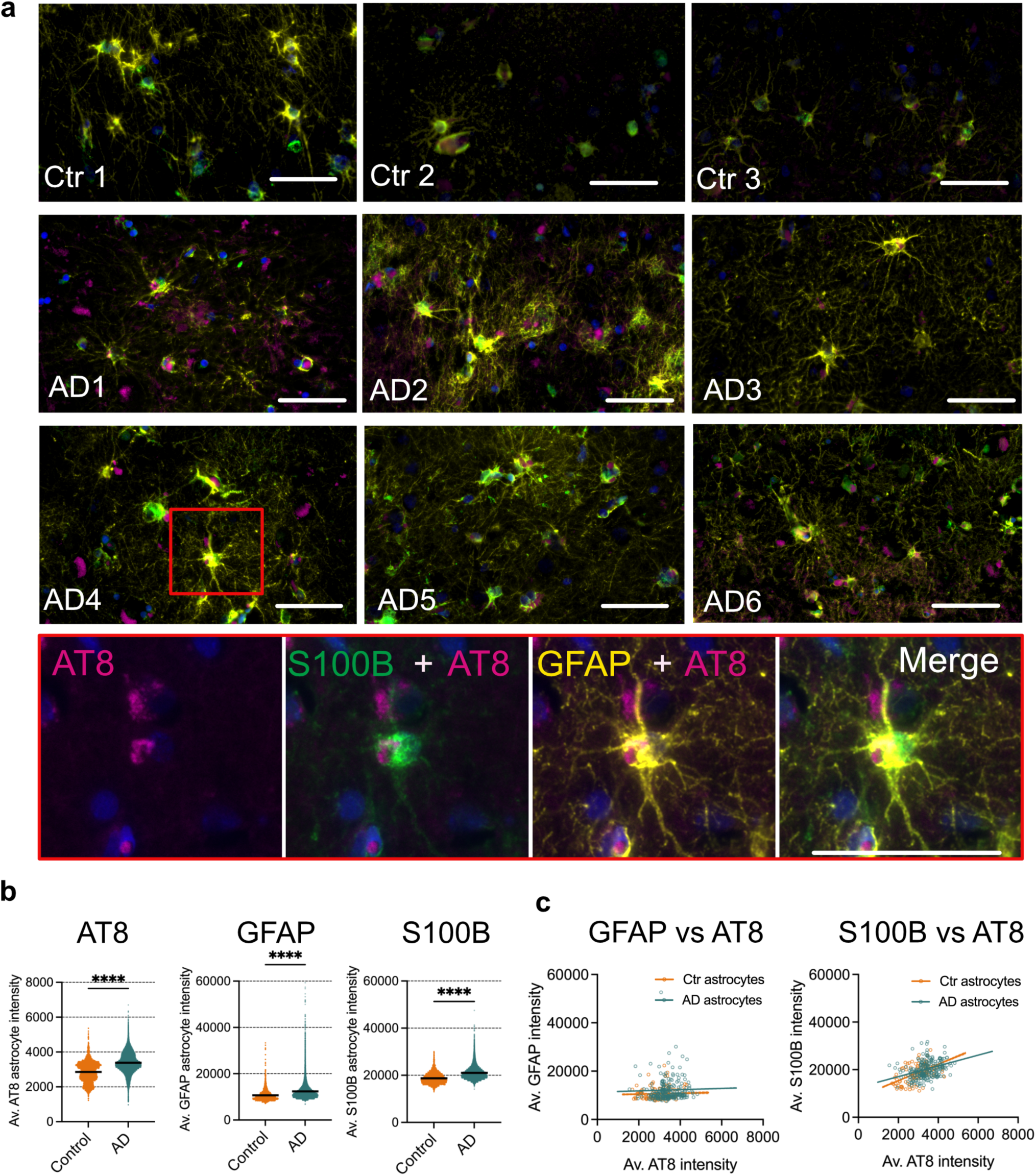
Characterization of human post-mortem temporal cortex shows tau inclusions associated with astrocytes. (a) Representative immunolabelling of astrocytes with antibodies against GFAP (yellow), S100b (green) and AT8 (purple) in temporal cortex tissue sections from 6 AD cases and 3 control brains. Lower panel shows higher magnification image of area indicated by the red box in AD4. Scale bar = 50 µm. (b) Scatter plots of AT8, GFAP and S100B intensity in individual astrocytes from AD (n=6) and Control (n=3) brain sections. Black lines represent mean intensities. (c) Pearson correlation analysis of intensity of AT8 immunolabelling relative to GFAP or S100B in individual astrocytes in temporal cortex of AD (n=6) and control sections (n=3). Data in (b) and (c) were analyzed from cells positive for both GFAP and S100B in grey matter of temporal cortex, for a combined total of 68510 astrocytes in AD tissue sections and 3217 astrocytes in control tissue sections using an unpaired t-test in (a) and Pearson correlation analysis in (b), ****p < 0.0001.

Astrocytes in AD brain are typically characterized by high levels of GFAP (Serrano-Pozo et al., 2011), as found here (Fig. 1b). S100b is a calcium binding protein expressed by astrocytes that was recently shown to hinder tau aggregation and seeding (Moreira et al., 2021). S100b immunoreactivity (Fig. 1b) was significantly higher in AD tissues relative to controls (F (8, 71722) = 5096, p<0.0001, R² = 0.3624), as with GFAP (F (8, 71723) = 232.7, p<0.0001, R^2^ = 0.02530), with significant variation between cases (Fig. S1b).

Pearson correlation analysis showed a neutral correlation between AT8 and GFAP intensity in both AD (r=0.032) and Ctr samples (r=0.035), whereas there was a positive correlation between AT8 and S100B for both Ctr (r=0.554) and AD (r=0.338) (Fig. 1c). Again, there was variation between individual cases, with correlation between AT8 and S100B ranging from positive (Ctrl 1, Ctrl 3, AD1, AD4, AD5) to none (Ctrl 2, AD6) (Fig. S1c). Similarly, some cases showed weak positive correlation between GFAP and AT8 levels (Ctrl 3, AD1, AD2, AD5), with others showing no correlation (Ctrl 1, Ctrl 2) or weak negative correlation (AD3, AD4, AD6) (Fig. S1c, d). Collectively, these findings highlight a complex and variable astrocytic reaction to tau pathology in AD brain.

### 2. Post-translational modifications of AD tau aggregates

Tau in AD brain shows several modifications according to disease stage (Wesseling et al., 2020), with significant variation in tau “strain” (Sanders et al., 2014) having consequences for tau seeding ability *in vitro,* as well as clinical outcomes (Dujardin et al., 2020; Haj-Yahya et al., 2020). Here, liquid chromatography-tandem mass spectrometry (LC-TMS) was used to identify sites of tau modification in sarkosyl-insoluble fractions from the AD and control brain samples used in this study. Two control cases (Braak stage 0-I) showed a small number of phosphorylated residues in the proline-rich domain (PRD) (Thr181, Ser202), which are associated with the early stages of tau aggregation (Wesseling et al., 2020), while any modifications present were below limits of detection for one control case. Phosphorylation at several residues were detected in the AD samples, particularly in the proline rich domain of tau, and to a lesser extent in the microtubule binding domain. All AD cases showed phosphorylation at some sites in the C terminal domain (Fig. 2a). The number of sites at which phosphorylation was detected and the extent of phosphorylation varied between AD case. A hierarchical cluster analysis based on the presence or absence of phosphorylation at specific sites yielded three groups. Controls 1-3, AD3 & AD5, and AD1,2 4 & 6 (Fig. 2b). AD3 and AD5, which were the only cases phosphorylated at Ser113 and Thr175/181 and S289, were placed in a distinct cluster from the other AD and control cases. Tau in AD6 did not show any phosphorylation between S185 and T214, whereas phosphorylation in this region was evident in the other five AD cases. All AD cases, but not controls, showed phosphorylation at S394/400/404 and S262, hallmark phosphorylation sites in AD (Wesseling et al., 2020) and S262 is linked with high tau seeding efficiency and disease worsening (Dujardin et al., 2020).

**Figure 2.**
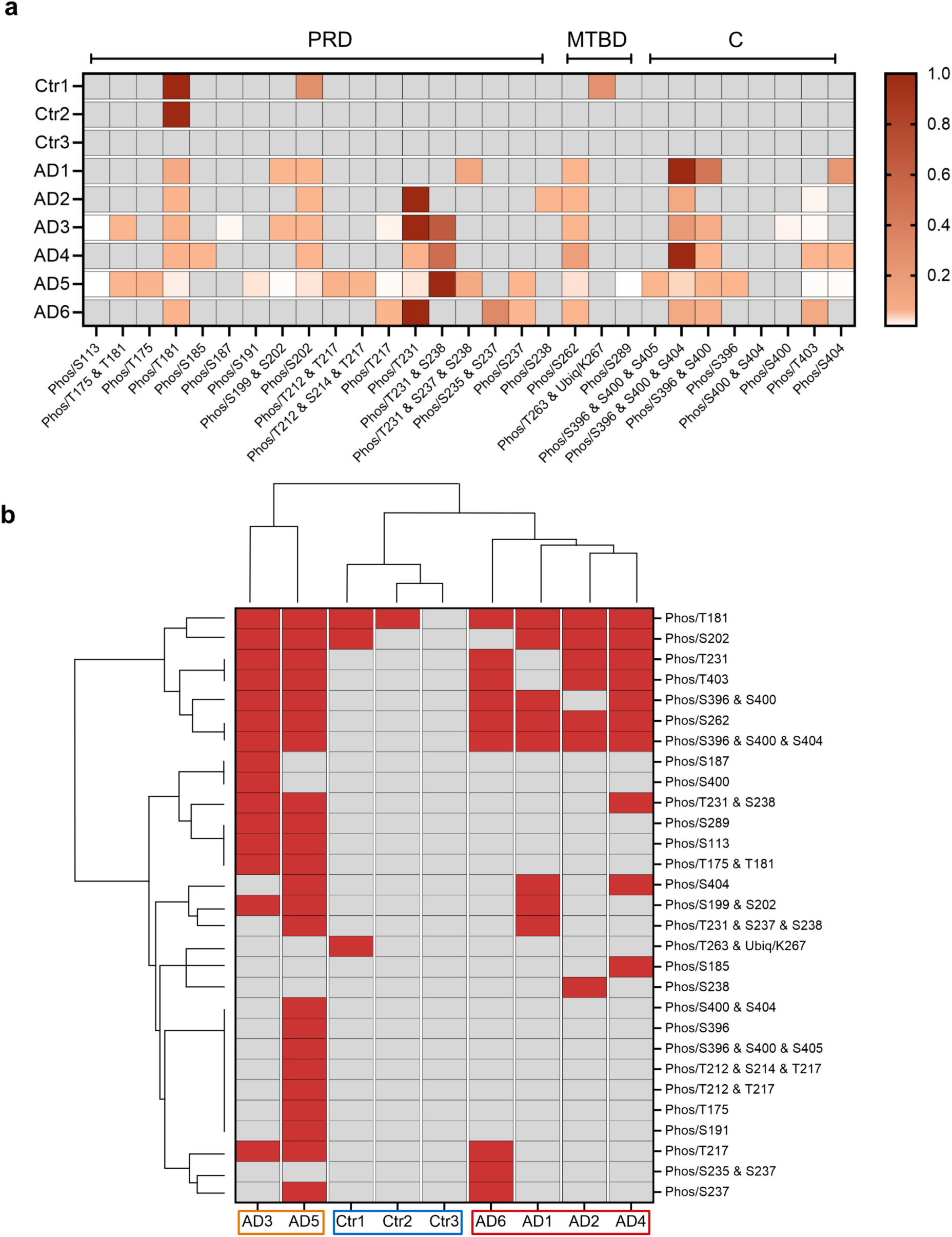
Characterization of sarkosyl-insoluble tau from postmortem human brain by LC-MS/MS. (a) Heatmap of tau phosphorylation sites within individual AD and control brain samples, quantified by the ratio of modified (phosphorylated) to equivalent unmodified peptides and normalized between 0 (white) and 1 (dark red) for each sample (grey = undetected) to show the relative abundance of tau phosphorylation sites within that sample. (b) The presence and absence of tau phosphorylation sites were used for unbiased hierarchical clustering of AD and control cases that determined 3 main groups: Ctr1-3; AD3,5; AD1,2,4,6. PRD=Proline rich domain; MTBD = microtubule binding domain; C= C terminus.

Peptide fragments from several other proteins were also detected in the sarkosyl-insoluble fraction (Fig. S2). An unsupervised hierarchical cluster analysis of the highest abundance of proteins separated control from AD cases (Fig. S2a). Within the AD cluster, AD3 and AD5 again appeared most distinct from AD2, AD4 and AD6 and the control cases. In addition to highest levels of tau, proteins such as Trypsin-1, RNA-splicing ligase RtcB homolog, and Ubiquitin-40S ribosomal protein S27a were highest in AD3/5. These proteins are involved in proteolytic activity, RNA processing, and protein degradation, suggesting distinct molecular alterations in these AD cases. Combined, 55 proteins were detected that differed significantly between AD and control tissue (Fig. S2b). The top most significantly enriched in AD brain included proteins involved in RNA processing such as U1 small nuclear ribonucleoprotein 70kDa (U1-70K), known to co-aggregate with AD tau (Bai et al., 2013; Bishof et al., 2018) and various small nuclear ribonucleoprotein-associated proteins (snRNPs), notable because RNA dysregulation may play a role in the progression of AD in some patients (Tijms et al., 2024). Other proteins significantly upregulated in AD cases included commonly associated AD proteins such as tau, apolipoprotein E (APOE) and amyloid-beta precursor proteins. Several proteins associated with the complement system were also detected such as complement C3 and complement C4-A. Agrin, a heparan sulfate proteoglycan seen to accumulate in AD brain was also highly upregulated in these aggregated tau fractions (Verbeek et al., 1999). It is possible that some of these proteins may participate actively in disease development if co-factors contribute to tau aggregation, seeding and/or spread.

### 3. Human astrocytes internalize AD brain derived tau aggregates

Recent data shows that human iPSC-derived astrocytes internalize neuronal debris (Mothes et al., 2023) and it might be reasoned that this accounts for tau immunolabelling in human PM brain astrocytes. However, growing evidence indicates that astrocytes can internalize different forms of human tau in an active process that might contribute to the spread of tau aggregates across diseased brain (Eisenbaum et al., 2023; Eltom et al., 2024; Martini-Stoica et al., 2018; Perea et al., 2019) .

Tau uptake and/or seeding is facilitated by the presence of similar forms of tau (Goedert et al., 2017; Wegmann et al., 2016) and single cell analysis of human brain shows that astrocytes express *MAPT* (Karlsson et al., 2021; Zhang et al., 2016). Here, astrocytes were differentiated from control iPSCs using a well-characterized protocol (Tcw et al., 2017). 70-days after differentiation, these cells express robust levels of mature astrocyte genes (Fig. S3a) and tau protein (Fig. S3b-d). As specific tau isoforms such as those expressed in mature brain may be more prone to misfolding and recruitment to facilitate tau spread (Chen et al., 2019; Zhong et al., 2012), we examined the expression of *MAPT* mRNA against alternatively spliced regions of tau using primer sequences that target exons 2 and 3 which encode the alternatively spliced N-terminal subunits of tau (0N, 1N, 2N tau) and exon 10 for the second microtubule binding domain (3R/4R) (Fig. S3b). In iPSCs and NPCs, 0N and 3R tau isoforms were the predominant source of *MAPT* mRNA. NPCs showed higher total levels of *MAPT* expression compared to iPSCs. As NPCs were differentiated into astrocytes, the proportion of 1N and 4R tau mRNA increased, and relatively low levels of 2N *MAPT* mRNA emerged during astrocyte maturation. These data suggest that astrocytes derived from iPSCs recapitulate developmentally regulated tau splicing, as observed at a slower rate, in iPS-neurons (Seto-Salvia et al., 2022).

We next spiked the media of 70-day iPSC-derived astrocytes with tau aggregates from the AD and control cases, and tau uptake by astrocytes was measured over 7 days (Fig. 3a). Immunolabelling of cells using the AT8 antibody allowed us to distinguish between endogenously expressed astrocytic tau (AT8 negative at baseline) and exogenously applied human tau (Fig. 3c). We found that astrocytes internalize tau in a time-dependent manner, showing greater volumes of AT8-positive tau aggregates after 7 days of incubation (Fig. 3a). Although tau aggregates from each AD case were internalized, we observed varying rates of tau uptake depending on the AD case used, with the slowest rates of uptake observed with tau from cases AD1, AD3 and AD5. Next, we studied the longer-term dynamics of tau handling by astrocytes. Here, tau-containing medium was removed after 7-days and iPSC-astrocytes grown in standard media for a further 14 or 28 days. Analysis of AT8 content after 14 days showed reduced AT8 volume suggesting that astrocytes degraded or otherwise cleared tau aggregates following the removal of tau from culture media, with the exception of AD3 (Fig. 3b). Examination of AT8 levels 28 days following tau removal from media showed increased AT8 reactivity suggesting that there was seeding of endogenous astrocyte tau in cases treated with tau aggregates from cases AD1, AD2, AD4 and AD6 (Fig. 3b, d). This feature was not observed with tau from cases AD3 and AD5 which showed further reductions in AT8 intensity, in addition to unique phosphorylation events (pSer113, pThr175/181, pS289) relative to the other samples.

**Figure 3.**
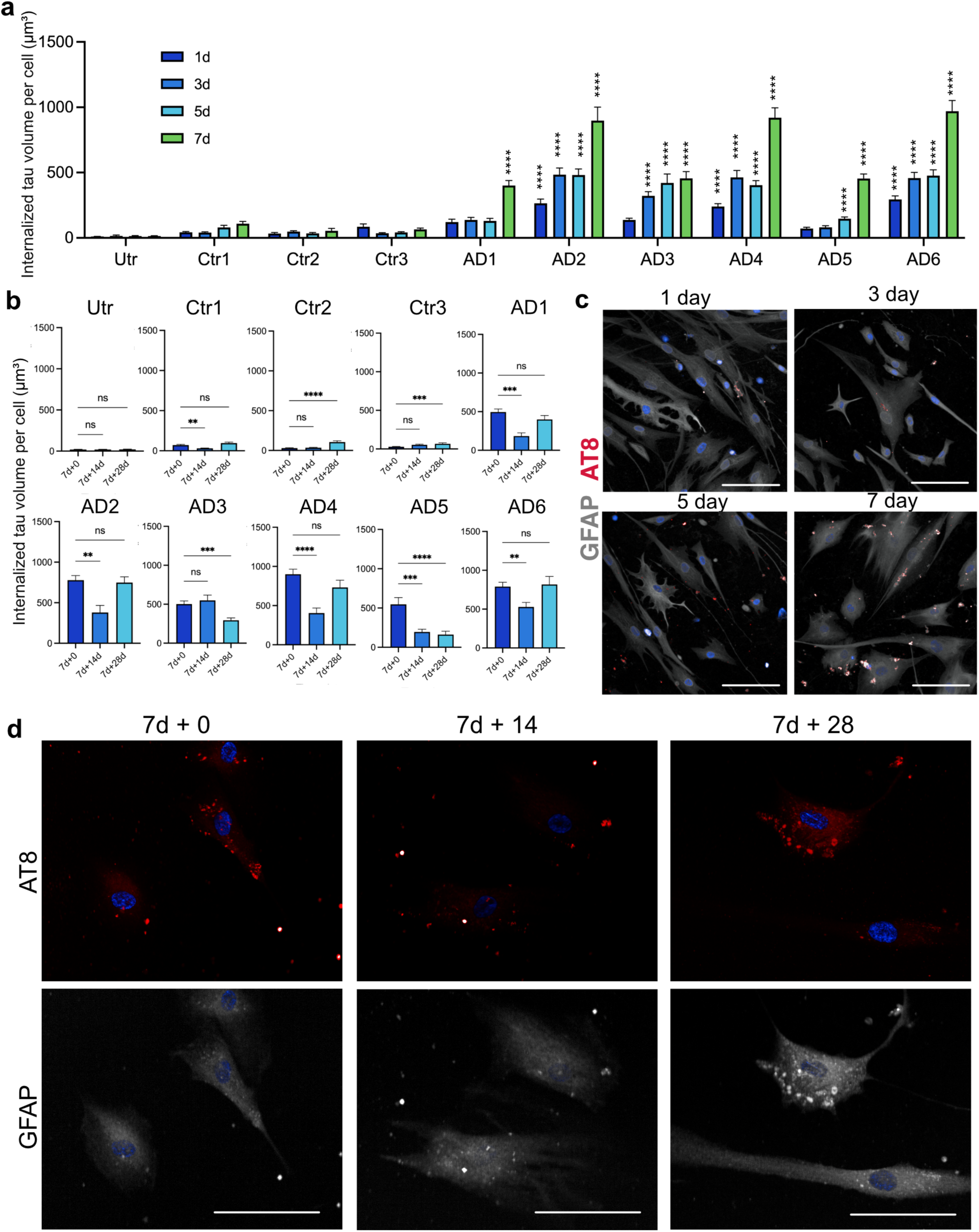
Astrocyte uptake of tau aggregates derived from AD postmortem tissue. iPSC-astrocytes were incubated with 35 ng/mL of sarkosyl-insoluble tau derived from post-mortem tissue of six AD cases and three equivalent control brain fractions. (a) Average detected volume of internalized AT8 positive tau aggregates in astrocytes after 1, 3, 5 or 7 days of exposure to sarkosyl-insoluble tau fractions. (n=171-419 cells per treatment condition across 3 experiments). (b) Average detected volume of internalized AT8 tau after 7-day tau incubation (+0), and at +14 and +28 days after tau removal from media. (n=400-500 cells across 3 experiments). (c) Representative immunolabelling of AT8 positive tau (red) internalized within GFAP positive (grey) astrocytes at 1,3,5 and 7 days after exposure to AD1 tau. (d) Representative immunolabelling of AT8 positive tau (red) internalized within GFAP positive (grey) astrocytes after 7d treatment (+0) and 14 days (7d +14) or 28 days (7d+28) after tau removal. Data is from three independent differentiations of iPS-astrocytes. Data is mean ± SEM. White scale bars = 100 µm. Statistical analysis by two-way ANOVA with Tukey’s multiple comparisons test to untreated cells in (a) and one-way ANOVA with Dunnett’s multiple comparisons test to baseline (7d+0) in (b). **** p<0.0001, *** p<0.001, ** p<0.01, * p<0.05, ns =not significant.

### 4. Tau aggregates disrupt astrocytic GFAP and S100b localization

After 7 days of exposure to human AD brain tau aggregates, immunolabelling showed significantly increased average S100b immunoreactivity following exposure to sarkosyl-insoluble fractions, particularly from AD cases (Fig. 4a). While variable, S100b was increased by exposure to all AD and control samples except for Ctr3 (Fig. 4b). Similar effects were observed for GFAP (Fig. 4a), where 4 of 6 AD cases showed significantly higher local GFAP immunoreactivity compared to only 1 of 3 control cases (Fig. 4b). However, we saw no significant global changes in *GFAP* or *S100B* gene expression relative to controls (Fig. S4b).

**Figure 4.**
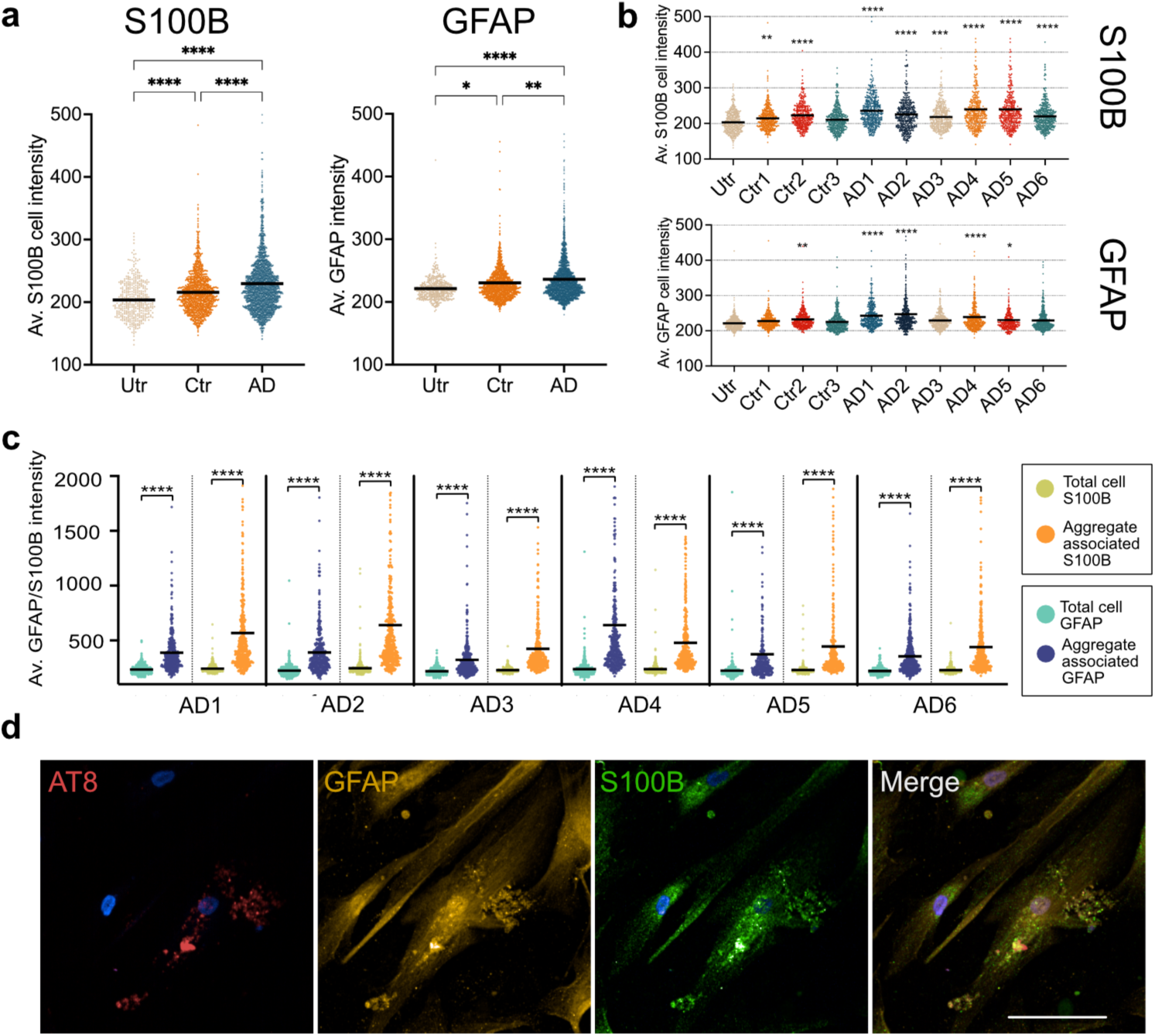
GFAP and S100B localize to internalized tau aggregates. Immunofluorescence intensity of GFAP and S100B in iPSC-astrocytes were measured after incubation with 35 ng/mL of sarkosyl-insoluble AD and control fractions for 7 days, (a) Scatter plots of average GFAP and S100B immunofluorescence intensity for combined AD treated (n=6), control treated (n=6) and untreated iPSC-astrocytes (2461, 1275 and 427 astrocytes respectively per treatment group across 3 experiments). (b) Scatter plots of average S100b and GFAP immunofluorescence intensity following exposure to tau from individual AD and control sarkosyl-insoluble fractions, relative to untreated iPSC-astrocytes (n=385-488 cells per treatment across 3 experiments). (c) Scatter plots comparing mean ‘total cell intensity’ relative to mean ‘aggregate-associated’ tau intensity after internalization of AD sarkosyl-insoluble tau aggregates (n=398-488 cells across 3 experiments). (d) Representative immunolabelling showing GFAP (yellow) and S100B (green) localizing at high levels around internalized AT8-positive tau aggregates (red) in astrocytes exposed to sarkosyl-insoluble AD1 tau for 7 days. White scale bar = 50 µm. Data is from three independent differentiations of iPS-astrocytes. Black bar is mean of individual cell data. Statistical analysis by one-way ANOVA with Dunnett’s multiple comparisons test to untreated in (b), and paired t test for GFAP/S100B total cell average vs aggregate-associated immunofluorescence in (c). **** p<0.0001, *** p<0.001, ** p<0.01, * p<0.05.

Pearson correlation analysis revealed significant associations between AT8 and both S100B and GFAP in astrocytes spiked with tau (Fig. S4a). These correlations are significantly more robust than were found in the AD brain analysis, indicating that there are confounding factors other than tau uptake that can affect tau association with astrocytes in diseased human brain.

There was a pronounced accumulation of both GFAP and S100b proximal to AT8+ tau within iPSC-astrocytes treated with tau from AD cases (Fig. 4c, d) that persisted following tau removal from media (Fig. S4c). These may suggest that GFAP and S100B are sequestered by tau inclusions, irrespective of whether seeding of endogenous tau was observed.

### 5. Internalization of tau aggregates from AD brain alters astrocytic gene expression

Astrocytes become reactive in AD, showing significant alterations in their transcriptional regulation (Escartin et al., 2021). To delineate broad gene expression changes common to the uptake of disease associated tau aggregates, bulk RNA-seq data from astrocytes treated with tau from AD cases or control brain extracts were grouped and compared to untreated astrocytes. Treatment with AD brain tau resulted in 96 significant (p<0.05) differentially expressed genes (DEGs), while treatment with equivalent control brain fractions showed 68 DEGs, of which 31 genes were common to both AD and control treated astrocytes (Fig. 5a, b). Of the overlapping genes, the correlation of fold change between both groups was very high (r=0.992), indicating these genes are dysregulated in a similar manner. These likely reflect responses to other components of the sarkosyl-insoluble fractions used. Cells treated with control brain extracts showed a higher number of downregulated genes than was observed following spiking with AD tau, and the mean fold change for both up and downregulated genes was highest in the AD group.

**Figure 5.**
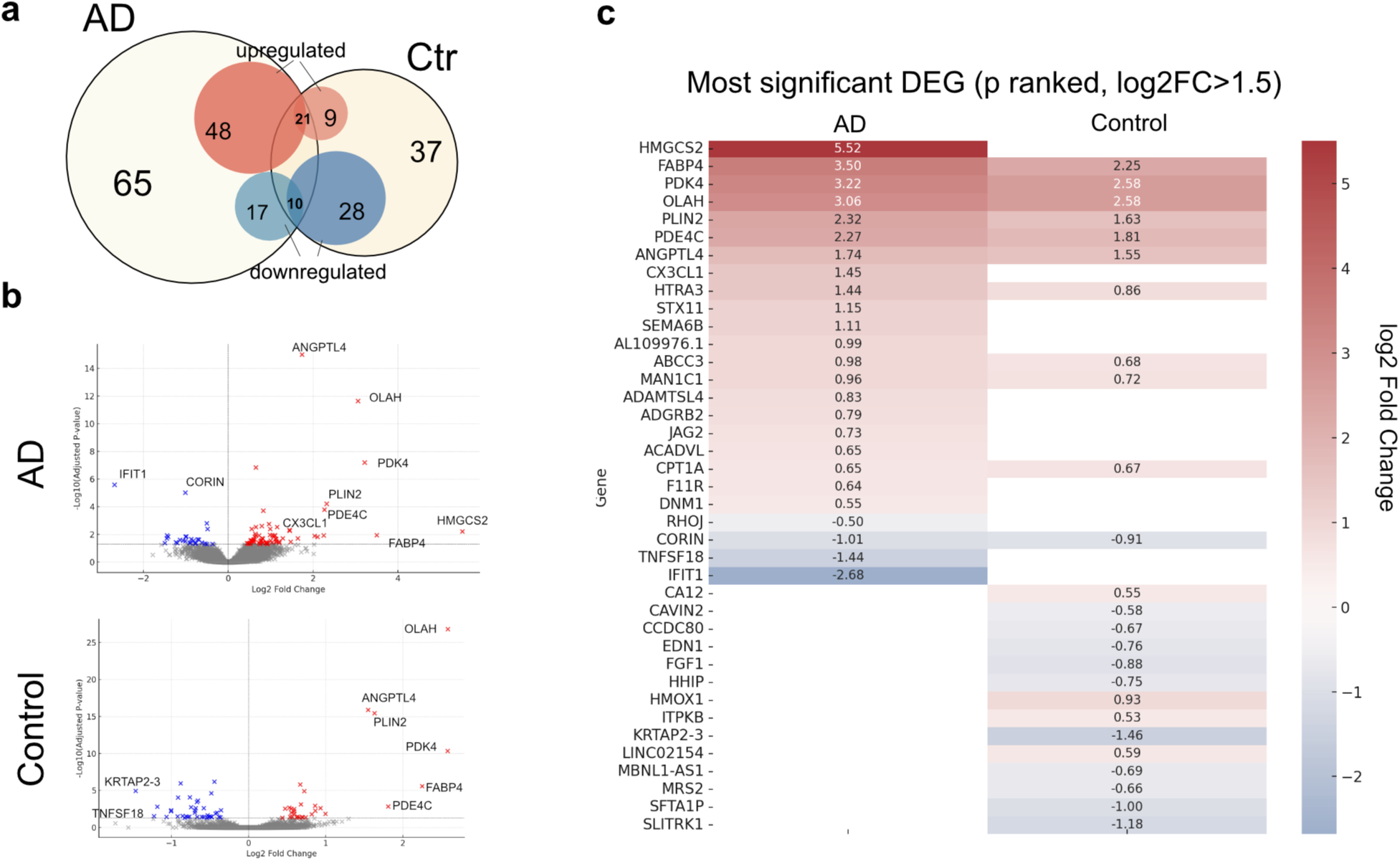
Differential gene expression in astrocytes after AD tau exposure. iPSC-astrocytes were treated with 35 ng/mL of tau from sarkosyl-insoluble fraction of (AD1-6) and control (Ctr1-3) brains for 7 days and compared to untreated astrocytes after bulk RNA-sequencing. Data was pooled to compare AD treated (n=6) and control treated (n=3) gene expression changes to untreated astrocytes across 3 experimental repeats. (a) Venn diagram of significantly (p<0.05) up (red) and down (blue) regulated genes in AD treated (left), control treated (right) and overlapping genes, relative to untreated astrocytes. (b) Volcano plot of significant upregulated (red) and downregulated (blue) DEGs (p<0.05) in AD and control treated astrocytes vs untreated controls, with annotation of the most significantly altered genes (p<0.01). (c) Top DEGs in both control and AD groups ranked by lowest p values and filtered by log2FC>1.5. Red gradient represents strength of upregulation and blue represents downregulation as per log2(Fold change) relative to untreated astrocytes.

The top 25 most significant DEGs (log2FC>1.5) show several unique and significantly altered genes in astrocytes exposed to AD tau, with known roles in astrocyte signaling and the clearance of pathological protein aggregates. For example, chemokine *CX3CL1 (*2.73-fold upregulation; p<0.01), regulates neuron-glial interactions in neurodegenerative diseases (Bivona et al., 2023; Subbarayan et al., 2022) and is upregulated in astrocytes in disease (Lindia et al., 2005), *IFIT1* (Interferon-Induced Protein With Tetratricopeptide Repeats 1) mediates astrocyte responses to viral infections (Zhang et al., 2017) and Tumor Necrosis Factor (Ligand) Superfamily Member 18 (*TNFSF18*) was downregulated 6.4-fold in the same treatment conditions. *HMGCS2* (mitochondrial 3-hydroxy-3-methylglutaryl-COA synthase 2), an enzyme that plays a key role in ketogenesis (Puchalska & Crawford, 2017), and is important for autophagic degradation of amyloid-beta precursor proteins (Hu et al., 2017) and tau (Hu et al., 2023) showed a large (45.9-fold) upregulation, perhaps indicating this as a key pathway in astrocytic tau processing (Fig. 5C).

We observed significant disparity in the number of DEGs across treatment conditions (Fig. 6a, Fig. S5a). Control conditions (Ctr1-3) exhibited relatively low numbers of DEGs, with counts of 8, 5, and 15, respectively. In contrast, AD-treated conditions showed a wider range of DEG counts, from as few as 2 in AD5 to as many as 581 in AD1, indicating a substantial variation in gene expression responses to spiking of cells with AD tau aggregates, which may relate to specific post-translational modifications of tau. When assessing the similarity between treatment groups using the Jaccard Index, the average similarity score among all AD treatment groups was approximately 0.130, while the control groups exhibited a higher average similarity score of approximately 0.333, indicating a more consistent gene expression response across control treatments. A hierarchical cluster analysis based on fold change of significant DEGs grouped AD1 and AD2 as unique from other treatment conditions (Fig. 6b). Of the remaining AD treatments, AD6 and AD4 showing greatest similarity, while AD3 and AD5, the two AD cases that were unable to seed astrocytic tau in our long-term assay (Fig. 3b), clustered more closely with controls (Ctr1-3).

**Figure 6.**
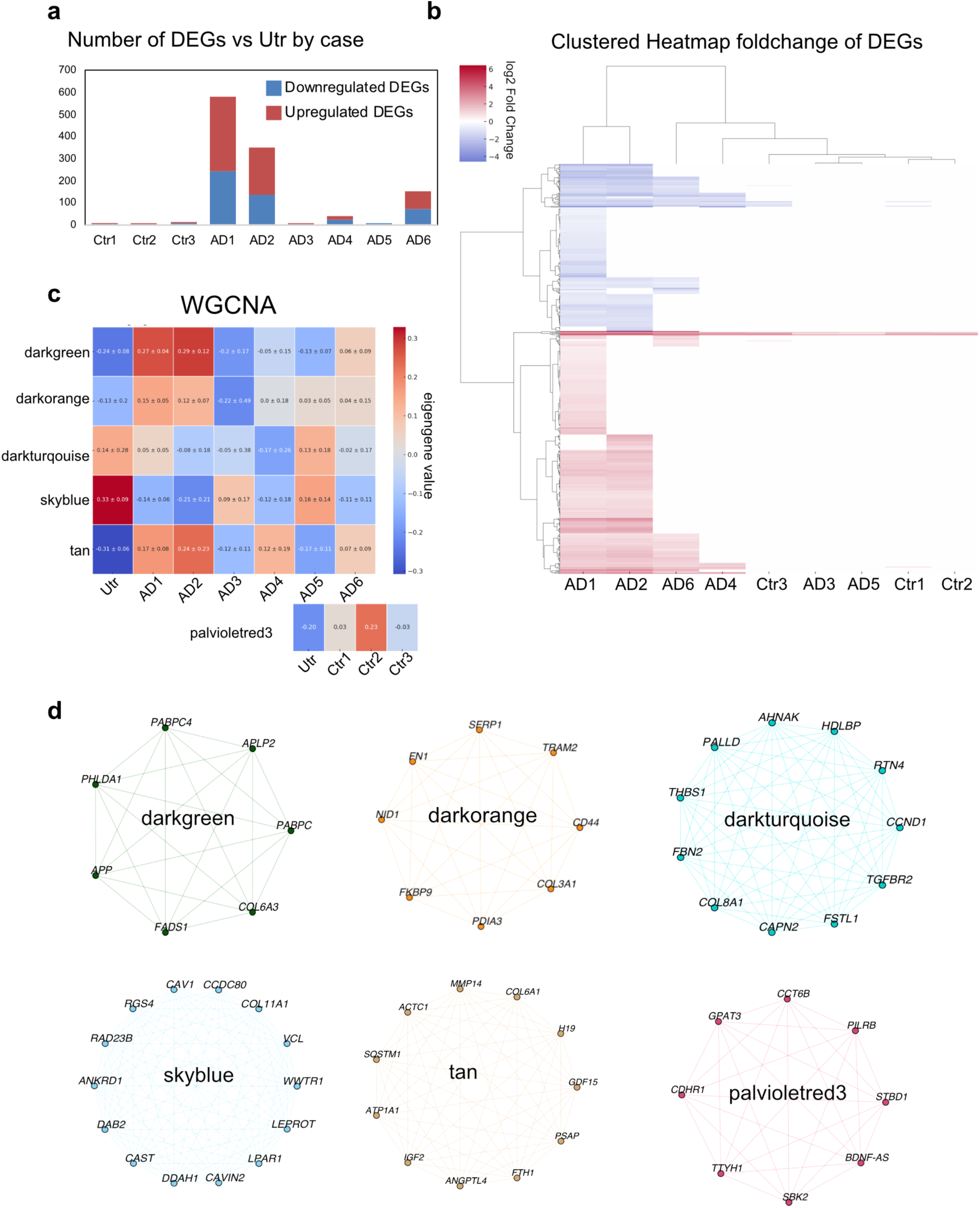
Heterogeneity in astrocytic gene expression response between AD cases. iPSC-astrocytes were treated for 7 days with 35 ng/mL of tau in sarkosyl-insoluble fractions of AD cases (AD1-6) and equivalent volumes of control brains (Ctr1-3) and compared individually to untreated astrocytes after bulk RNA-sequencing. (a) Total number of up- (red) and down-(blue) regulated DEGs in astrocytes exposed to samples from individual AD and control cases, compared to untreated cells. (b) Hierarchical clustered heatmap showing all significant genes (p<0.05) for astrocytes treated with each case compared to untreated. (c) Weighted correlation network analysis (WGCNA) heatmap depicting modules with most consistent expression changes across technical repeats, compared to untreated astrocytes. The strength of gene expression for each module is represented by its eigengene value representing trend of upregulation (red) or downregulation (blue) of gene in that module compared to untreated astrocytes. Each module displays a mean of 3 technical repeats ± SD. (d) Highest expressed genes for each module from (c) depicted in network diagrams.

Weighted gene co-expression analysis (WGCNA) identified 5 modules across AD tau-treated cells and one control module that were selected based on uniformity of eigengene values across technical replicates relative to untreated cells (Fig. 6c). The highest expressed genes in each module are highlighted in network maps (Fig. 6d) and expression heatmaps (Fig. S5b).

The **darkorange** module contains genes involved in extracellular matrix interactions (*NID1, FN1* and *COL3A1*) (Lau et al., 2013; Soles et al., 2023) protein folding and ER stress response (*PDIA3, FKBP9, TRAM2*) (Ajoolabady et al., 2022; Blair et al., 2015; Cassano et al., 2023), as well as *CD44*, which may have a role in mediating neuroinflammatory responses (Sawada et al., 2020). AD1 and AD2 demonstrated the strongest deviation from untreated cells and had the highest eigengene values, whereas AD3 was similar to untreated cells, again indicating variability in the response to tau from different AD cases.

The **darkgreen** module was starkly upregulated in AD1 and AD2 compared to untreated cells, with only moderate differences between cells treated with tau from the other AD cases and untreated controls. This module contains genes associated with amyloid precursor protein processing (*APP, APLP2*) (Zhang et al., 2011) and lipid metabolism (*FADS1*) (Mathias et al., 2014). It also contains genes with functions in mRNA stability and translation (*PABC1* and *PABPC4*) (Kini et al., 2014; Qi et al., 2022), and *PHLDA1* which is linked to microglia activation (Han et al., 2020).

The **tan** module contains genes involved in autophagy (*SQSTM1*) (Huang et al., 2023; Kageyama et al., 2021), lysosome function and lipid metabolism (PSAP) (He et al., 2023), extracellular matrix remodelling (*MMP14* and *COL6A1*) (Gregorio et al., 2018; Winkler et al., 2020), growth factors that influence cell growth, differentiation and survival (*IGF2)* (Alberini, 2023) whose expression has been noted to decrease in AD patients vs controls, and which has been touted as a potential therapeutic target (Fitzgerald et al., 2023). Other hits included *GDF15*, a mitochondrial stress response molecule (Chiariello et al., 2022) and *ANGPTL4* (Fernandez-Hernando & Suarez, 2020), iron metabolism (*FTH1*), which has been linked to ferroptosis in AD brain via single cell RNA-seq analysis (Dang et al., 2022), and ion transport across membranes (*ATP1A1*) (Biondo et al., 2021). Again, this upregulation was particularly pronounced when cells were treated with tau from AD1 and AD2, while cells treated with AD3 and AD5 extracts showed expression similar to untreated cells.

For the **darkturquoise** module, expression was reduced in the majority of AD treated astrocytes compared to untreated, except for AD5. This module again contains genes involved in extracellular matrix organization including THBS1 (Atanasova et al., 2019), *FBN2* (Frederic et al., 2009), and *COL8A1*, the latter being upregulated in an AD specific astrocyte cluster analysis (Sadick et al., 2022). Other hits include genes involved in growth factor signaling such as *FSTL1* and *TGFBR2* (Ghasempour et al., 2022; Jin et al., 2017). *RTN4* is within this module, a regulatory transmembrane protein implicated in neurodegenerative disease (Gns et al., 2022; Kulczynska-Przybik et al., 2021). *PALLD* encodes a cytoskeletal protein involved in cell shape and motility (Parast & Otey, 2000). Genes related to protein synthesis and trafficking (*HDLBP* (Zinnall et al., 2022) and *AHNAK* (Gentil et al., 2001) were also detected. *CCND1* also present in this module, is important for cell cycle regulation and has been identified as a transcriptional regulator of reactive astrocytes (Burda et al., 2022), as well as *CAPN2*, a protease studied in AD models and with therapeutic potential (Ajoolabady et al., 2022; Ono et al., 2016).

The **skyblue** module was also decreased when cells were treated with tau from most AD cases compared to untreated. It contains genes such as *CAV1, CAVIN2*, and *LPAR1,* important in cell signaling and membrane dynamics (Annabi et al., 2017; Jasmin et al., 2012; Xiao et al., 2021).The involvement of genes related to the extracellular matrix is again highlighted by *COL11A1* (Liu et al., 2021) and *VCL* (Mandal et al., 2021), reinforcing the importance of cell-matrix interactions. Additionally, *WWTR1* (Yu et al., 2021)and *RGS4* (Gns et al., 2022) underscore transcriptional regulation and signal transduction genes, with both previously noted as hub genes altered in AD progression. The module also showcases *DDAH1* and *CAST* for their contributions to protein metabolism (Averna et al., 2003; Hu et al., 2021), along with *DAB2* (Ogbu et al., 2021) and *RAD23B* (Jensen et al., 2018) which have roles in endocytosis and nucleotide excision repair. There was a general decrease in expression across all AD cases compared to untreated cells, most profoundly for those treated with AD1, AD2, AD4 and AD6 tau, but not AD3 and AD5, suggesting that these genes are altered with pathological tau seeding and accumulation.

Finally, the **palevioletred3** module highlights genes upregulated following exposure to control brain extracts relative to untreated cells. The genes in this module are involved in various cellular processes including glycophagy (*STBD1)* (Zhu et al., 2014), kinase activity (*SBK2)* (Manning et al., 2002), immune response regulation (*PILRB*) (Karch et al., 2016) lipid biosynthesis *GPAT3* (Fan et al., 2023), and *BDNF-AS* which regulates the expression of *BDNF* (Ghafouri-Fard et al., 2021).

These insights underscore the complex and heterogeneous astrocyte responses to AD tau, spotlighting the significant role of distinct gene modules in AD progression and the potential influence of these genes on astrocyte physiology I response to pathological tau processing.

## Discussion

Astrocytic tau accumulations are characteristic of several primary tauopathies including progressive supranuclear palsy and corticobasal degeneration. Recent studies (Nolan et al., 2019; Richetin et al., 2020) have also observed tau aggregates within astrocytes in AD brain, although their specific role in AD is not well understood. Consequently, several aspects require further exploration: the frequency and dynamics of astrocytic tau inclusions in AD brain, the capacity of astrocytes to eliminate these inclusions or contribute to tau spread, and the impact of such inclusions on astrocyte function. We make progress in illuminating some of these roles for astrocytes in AD using a relatively small number of AD cases. We show differences in astrocyte responses to tau from different AD cases which likely reflects molecular heterogeneity of tau, giving some insights into the nuance required for interpreting astrocyte responses to tau in disease.

Using high-throughput immunofluorescence screening of temporal cortex, we show that astrocytes are infrequently associated with pathological tau inclusions in the AD cases examined here, an observation which extended to one control case that showed early Braak staging, suggesting that astrocytes may develop tau inclusions early in disease. In AD brain, astrocytes come into contact with ‘ghost’ tangles that exist in the extracellular spaces (Perez-Nievas & Serrano-Pozo, 2018; Probst et al., 1982), and potentially internalize tau aggregates along with neuronal debris, as previously described (Mothes et al., 2023). However, astrocytes also internalize isolated tau (Eisenbaum et al., 2023; Eltom et al., 2024; Martini-Stoica et al., 2018; Perea et al., 2019). To further explore responses of astrocytes to pathological tau, we optimized a human cell-based assay for seeding with AD brain derived sarkosyl-insoluble tau aggregates, considering that synthetic tau fibrils, with their distinct conformational structure (Shi et al., 2021), might elicit different cellular responses. These fractions were found to contain several other protein components, and unbiased cluster analysis distinguished between samples that induce tau seeding in astrocytes (AD1, AD2, AD4, AD6) and those that do not (AD3, AD5), suggesting that co-factors as well as post-translational modifications, may influence tau seeding and potentially spread in diseased brain.

We found that astrocytes internalized tau aggregates at a relatively slow rate in comparison to microglia (Konishi et al., 2022; Loov et al., 2015). Interestingly, the rate of tau uptake varied depending on the AD case, as did the propensity to seed endogenous tau aggregation. One possible explanation is that, although AD tau fibrils share a common proteopathic core (Fitzpatrick et al., 2017), their post-translational modification profiles can significantly impact tau seeding (Dujardin et al., 2018; Dujardin et al., 2020). Notably AD3 and AD5, two cases we found to have slow rate of uptake and more efficient clearance by astrocytes, had unique phosphorylation profiles that distinguished them from other cases that we studied, including peptides phosphorylated at sites that have previously been shown to negatively correlate with tau seeding and disease progression (Dujardin et al., 2020). Moreover, the abundance of peptides modified at S262 was relatively low for AD5 compared to other cases, and phosphorylation at this site has shown positive correlation with tau seeding (Dujardin et al., 2020). Further, phosphosites S113 and S175, uniquely detected in AD3 and AD5, as well as S191 in AD5, were more common in a cluster of AD cases that had lower tau burden (Wesseling et al., 2020). While it has often been assumed that greater overall levels of phosphorylation promote tau aggregation, our work is in line with recent studies that show that tau aggregation is dependent on specific sequence modifications in tau (Dujardin et al., 2020; Kamath et al., 2021), and that some modified sites can in fact inhibit aggregation (Haj-Yahya et al., 2020). More work is required to carefully delineate how specific modification sites of tau can affect uptake, seeding and downstream cell type response.

Emerging evidence supports the role of astrocytes in not only internalizing tau but also facilitating its spread to neighboring cells, thus propagating tau pathology (Eltom et al., 2024; Mothes et al., 2023). Our work further suggests that tau modifications impact tau uptake and seeding in astrocytes that may contribute to this process. Our work agrees with others that human astrocytes express *MAPT* (Karlsson et al., 2021), and therefore provide a source of tau (albeit at low levels) for templated misfolding. Much as tau aggregation affects neuronal function, we found that tau accumulation in astrocytes affected astrocytic proteins such as GFAP and S100b, both of which are linked to AD progression (Middeldorp & Hol, 2011; Sheng et al., 1994; Van Eldik & Griffin, 1994). S100B localizes with tau in neuroblastoma cells and can prevent tau seeding and limit liquid-liquid phase separation *in vitro* (Moreira et al., 2021; Moreira & Gomes, 2023). However, S100B may also contribute to tau hyperphosphorylation through DKK-1 upregulation (Esposito et al., 2008). Its persistence around internalized tau aggregates after exogenous tau removal in our assay indicates an insufficient capacity to prevent endogenous astrocytic tau seeding. GFAP, indicative of astrocyte reactivity (Escartin et al., 2021), also showed a positive correlation with AT8 tau internalization, and may be sequestered by tau aggregates in the cytosol. Indeed, tau fibrils do not form as homogenous aggregates in AD brain, and several studies have shown other proteins are recruited into larger fibrils with tau (Rahman & Lendel, 2021), as also suggested by our analysis of sarkosyl-insoluble fractions.

Astrocyte reactivity has wide-ranging implications for AD progression, with unique signatures in AD brain (Habib et al., 2020). Responses to Aβ and tau pathology can differ (Jiwaji et al., 2022), and efforts are ongoing to resolve transcriptional changes during the spatial and temporal progression of AD pathology (Choi et al., 2023; Serrano-Pozo et al., 2022). Our RNA-seq analysis revealed highly significant DEGs unique to the AD group, including those related to the complement system (*CX3CL1*) and mitochondrial function and autophagic processes (*HGMCS2*), both previously implicated in AD pathology (Bivona et al., 2023; Hu et al., 2023; Hu et al., 2017; Subbarayan et al., 2022). While control brain extracts induced some overlapping gene expression which might reflect effects of other protein components in the sarkosyl-insoluble fraction, AD brain extracts produced a higher number of DEGs with a larger effect size. Again, variability in astrocyte gene changes was noted depending on the case used. Tau from AD3 and AD5, distinguished by their PTM profiles, showed slower uptake rates and did not seed further tau aggregation, and exhibited fewer DEGs. WGCNA analysis unveiled gene modules intersecting with cardinal features of AD pathology. The darkgreen module accentuated genes involved in amyloid precursor protein processing, including *APP* and *APLP2*, potentially linking tau uptake-induced astrocytic gene expression alterations to amyloid-beta pathology. This module also contained genes related to lipid metabolism, mRNA stability, and microglia activation.

We also observed changes to genes related to protein folding and endoplasm reticulum (ER) stress response after tau uptake, another mechanism implicated in AD onset and development (Ajoolabady et al., 2022). *PDIA3*, an ER isomerase expression is altered in 3xTg-AD mice (Cassano et al., 2023), and *FKBP9* part of a class of chaperones linked to AD progression (Blair et al., 2015), has been directly implicated in a prion seeding assay *in vitro* (Brown et al., 2014). In addition, *TRAM2* is a member of the translocon that is involved in the posttranslational processing of proteins at the ER membrane (Voigt et al., 1996).

Several modules demonstrated genes related to autophagy, lysosome and proteasome function, emphasizing the protein clearance pathways altered as astrocytes process tau aggregates. These included *HMGCS2*, *SQSTM1* (Huang et al., 2023), and *PSAP*, the latter being important for dopaminergic lipid homeostasis in a Parkinson’s model (He et al., 2023). Other modules linked genes related to protein metabolism such as *CAST* (calpastatin), *CAPN2* (calpain-2), *HDLBP* (Zinnall et al., 2022), and *AHNAK* (Gentil et al., 2001). Calpain activity has been shown to be upregulated prior to tau phosphorylation and loss of synaptic cells in AD brain (Kurbatskaya et al., 2016), and calpain-2 specifically can create tauopathy-associated tau fragments (Cicognola et al., 2020). Inhibition of calpastatin degradation reduced neuropathology in mouse models of Huntington’s (Hu et al., 2021), highlighting this pathway as a potential therapeutic target to reduce neuronal death.

Multiple modules contained genes integral to ECM organization and maintenance, including *MMP14*, *COL6A1*, *NID1*, *FN1*, *THBS1*, *FBN2*, *COL8A1*, *COL3A1*, *COL11A1*, and *VCL* (Atanasova et al., 2019; Frederic et al., 2009; Gregorio et al., 2018; Lau et al., 2013; Liu et al., 2021; Mandal et al., 2021; Soles et al., 2023; Winkler et al., 2020). This suggests astrocyte involvement in restructuring the brain’s extracellular environment in response to AD tau pathology. Components of the ECM are known to form part of amyloid structures in AD brain (Rahman & Lendel, 2021), and ECM remodeling can influence the distribution and aggregation of Aβ (Moretto et al., 2022). The rapid alterations in ECM related genes in our assay suggest that astrocytes play a role in this dysfunction.

Overall, our study reveals that astrocytes efficiently internalize AD associated tau aggregates but process them in different ways depending on characteristics of the tau itself. We also show that the expression of endogenous astrocytic tau may facilitate tau seeding in astrocytes during AD progression. We have identified that such uptake significantly influences astrocyte gene expression, pinpointing several genes previously linked with AD progression. This implies that the internalization of tau may prompt disease-associated astrocyte phenotypes, which likely disrupt interactions of astrocytes with other neural cell types. neurons and other glial cells. Together, this study underscores the significance of understanding the diverse effects of tau on astrocytes within patient cohorts, distinctions crucial for developing successful therapies targeting astrocytic functions.

## Materials and Methods

### 1. Key Resources Table

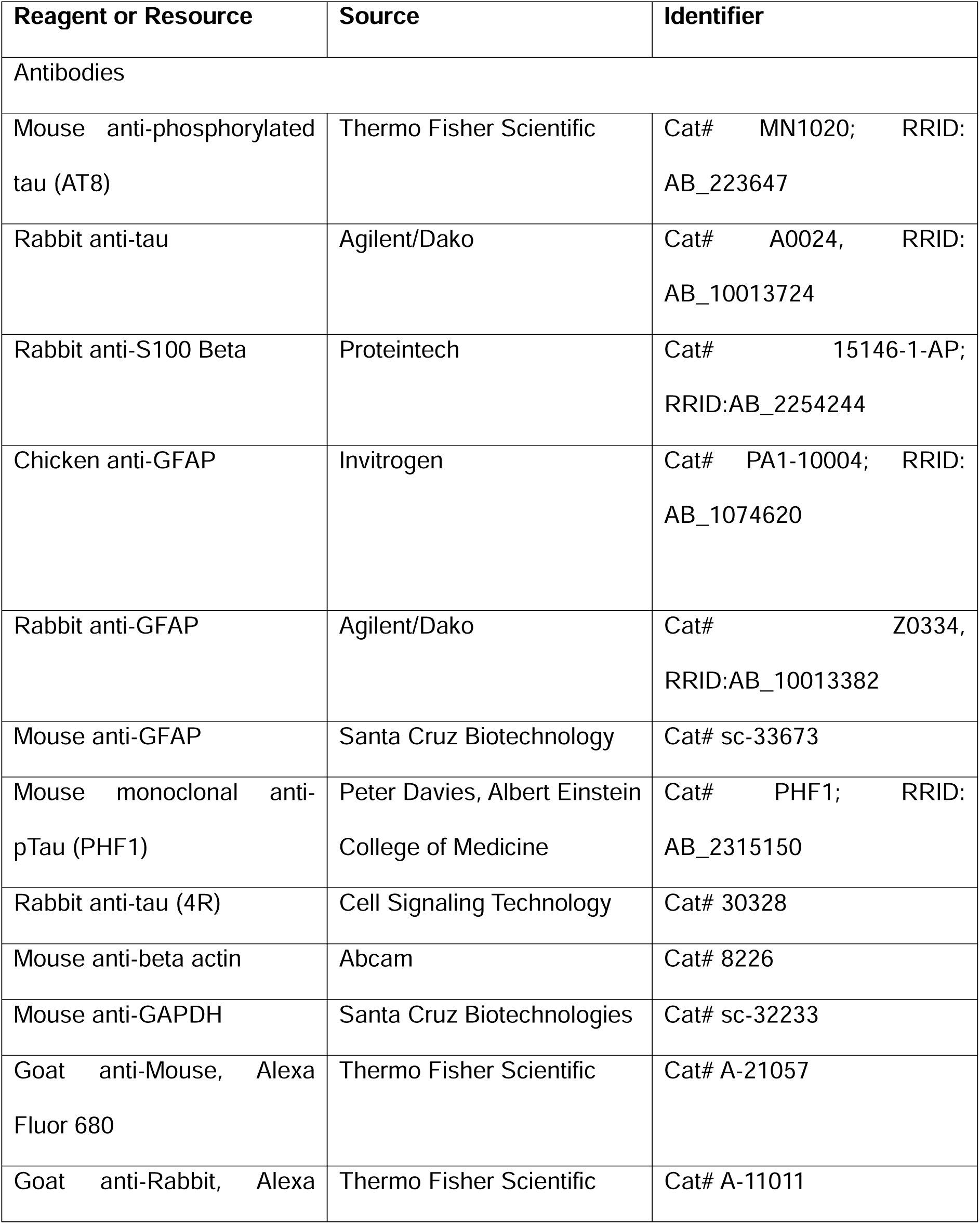
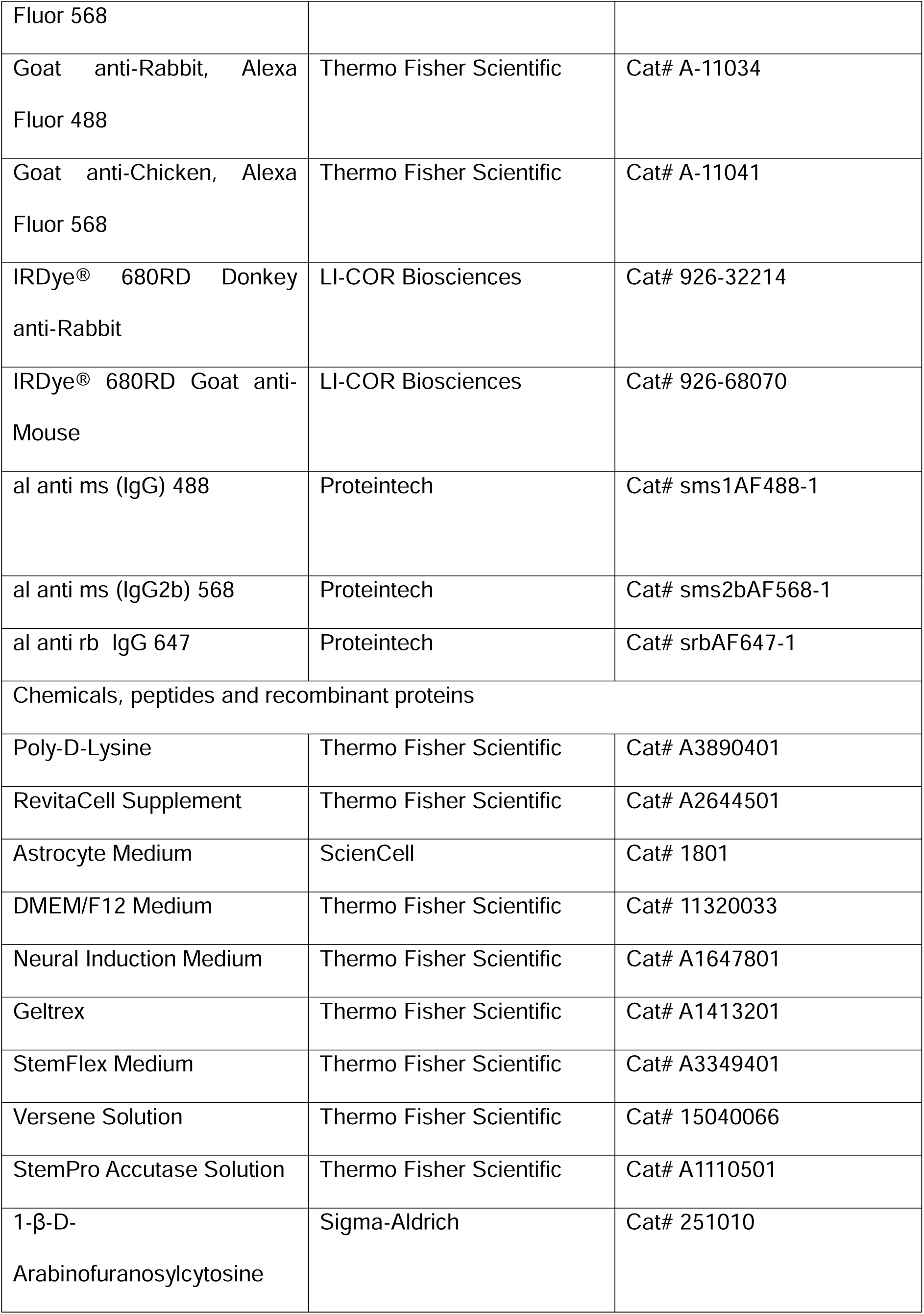
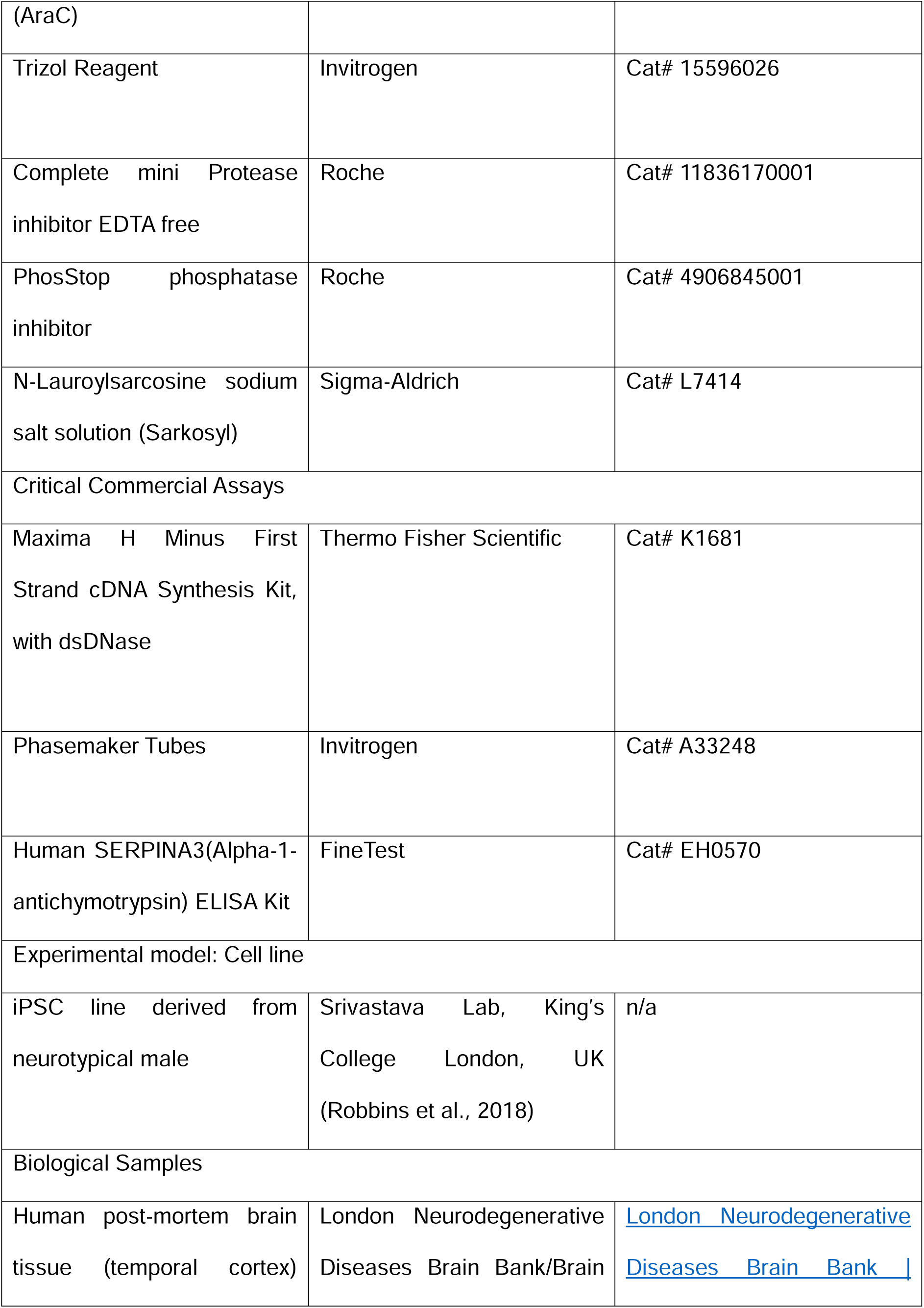
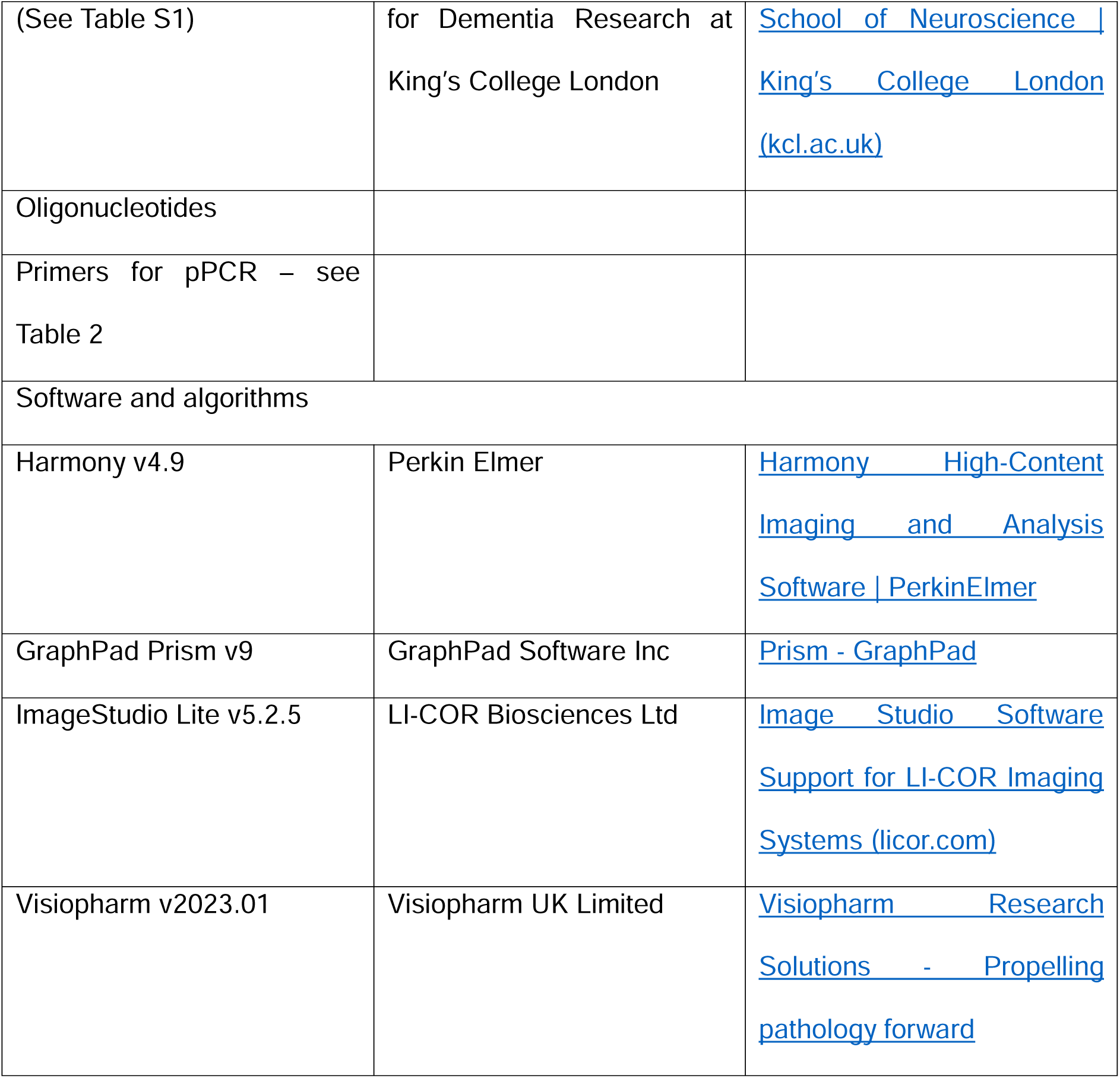

**Table S1.** Human brain samples utilized in this study with age at death, sex, post-mortem delay (PMD), pathological diagnosis and ApoE status (if known)

| Alias | Age | Sex | PMD | Pathological Diagnosis | ApoE |
| --- | --- | --- | --- | --- | --- |
| AD 1 | 81 | M | 74 | AD Braak VI |  |
| AD 2 | 88 | M | 46 | Alzheimer's disease: Definite Braak VI | 3/4 |
| AD 3 | 72 | M | 5 | AD, Braak VI with marked amyloid angiopathy |  |
| AD 4 | 97 | F | 12 | Alzheimer's disease Braak V | 2/3 |
| AD 5 | 72 | M | 5 | Alzheimer's disease Braak VI with marked amyloid angiopathy | 3/3 |
| AD 6 | 87 | F | 48 | Alzheimer's disease Braak VI with moderate amyloid angiopathy | 2/4 |
| Ctr 1 | 77 | M | 11 | Non-diseased |  |
| Ctr 2 | 86 | M | 6 | Non-diseased | 3/3 |
| Ctr 3 | 55 | F | 12 | Minimal tau pathology consistent with HP-tau stage I | 3/3 |

**Table 2.**
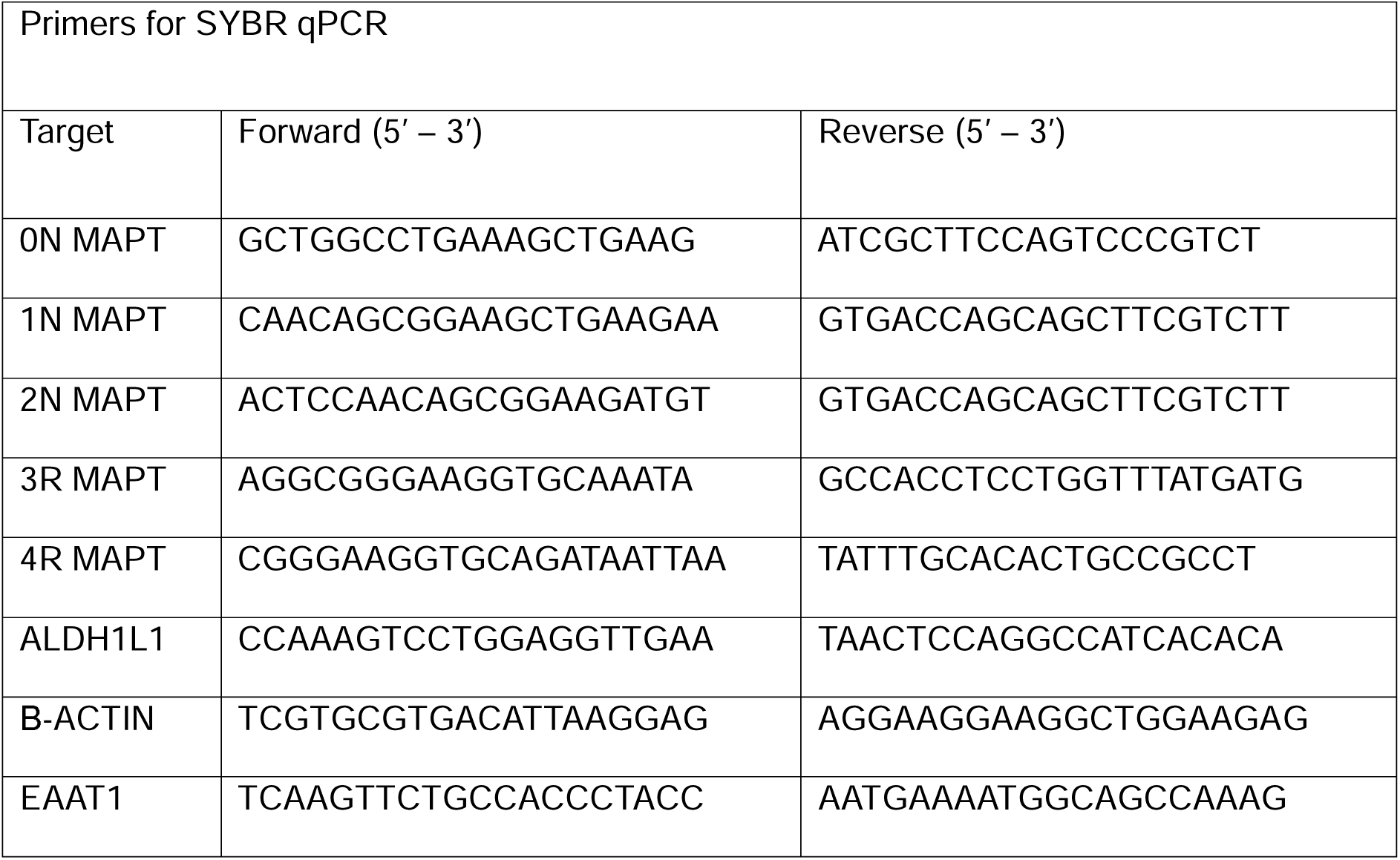
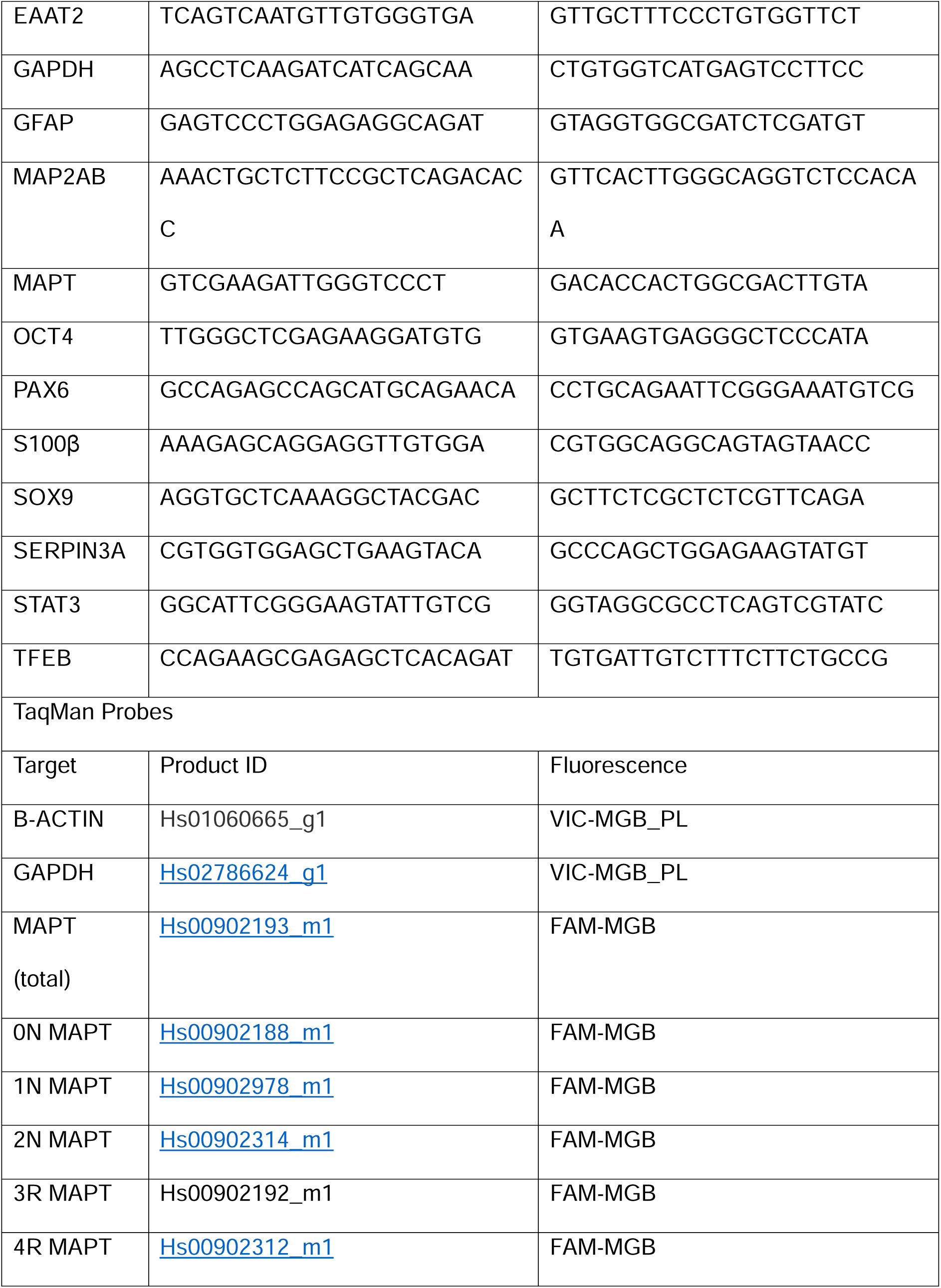
Primer sequences used for SYBR RT-qPCR and pre-designed TaqMan probes.

### 2. Resource Availability

#### Lead contact

Further information and requests for resources and reagents should be directed to and will be fulfilled by the Lead Contacts, Dr Matthew Reid or Prof Wendy Noble

#### Materials availability

No samples are available upon request.

#### Data and code availability

Any additional data in this paper for reanalysis is available upon request.

### 3. Experimental Model and Subject Details

#### Human iPSC line

The iPSC line used in this study (CTR_M3_36S) was received as a donation from Prof. Deepak Srivastava at King’s College London, previously generated from hair-root derived keratinocytes of a healthy control male, as described in (Cocks et al., 2013). The line was created with a CytoTune™-iPS 2.0 Sendai Reprogramming Kit (Cat# A16517; Invitrogen). This line was previously characterized for stable karyotype, differentiation into 3 germ layers, and efficient clearance of reprogramming transcription factors (Robbins et al., 2018).

Consent for storage of tissue and iPSCs and explicit consent for use in subsequent research was obtained by the original lab, following the guidelines of the UK Human Tissue Act 2004. Specific Research Ethics Committee approval is not a legal requirement for generating iPSCs from human donor tissue.

#### Human samples

Post-mortem human brain samples were requested from the London Neurodegenerative Diseases Brain Bank/Brains for Dementia Research at King’s College London. All human tissue collection and processing were carried out under the regulations and licensing of the Human Tissue Authority, and in accordance with the UK Human Tissue Act, 2004.

Samples were taken from the temporal cortex by an experienced pathologist. Pathological diagnosis was obtained by routine pathological analysis after donation to the brain bank and conducted by an experienced pathologist.

### 4. Method Details

#### Extraction and Quantification of Tau from Human Tissue

Tau was extracted from post-mortem human brain tissue using a protocol modified from Greenberg & Davies (1990). Briefly, brain tissue samples were homogenized at 100 mg/mL in Tris-Buffered Saline (TBS: 50 mM Tris-HCl, 150 mM NaCl, pH 7.4) containing 2 mM EGTA, 10% (w/v) sucrose, Complete Mini Protease Inhibitor Cocktail, and PhosSTOP phosphatase inhibitors (Roche, Basel, Switzerland). Homogenization was performed using a Tissue Master homogenizer (Omni International, USA). Sarkosyl (Sigma-Aldrich, St. Louis, MO, USA) was added to the homogenate to a final concentration of 1% (v/v), and samples were agitated at room temperature for 30 minutes. Samples were centrifuged at 136,000 x g for 1 hour at ambient temperature using a Beckman Coulter Optima MAX-XP Ultracentrifuge with a TLA-55 rotor (Beckman Coulter, CA, USA). The resulting sarkosyl-insoluble pellet was washed in homogenization buffer containing 1% sarkosyl, re-centrifuged, and then resuspended in sterile Dulbecco’s PBS (Thermo Fisher Scientific, MA, USA). The final sarkosyl-insoluble tau suspension was sonicated using a Bandelin Sonopuls HD 2070 (BANDELIN electronic GmbH & Co, Berlin, Germany). For quantification, tau was analyzed using SDS-PAGE and western blotting. Known concentrations of recombinant human tau (Human Tau Protein Ladder 6 isoforms; Sigma-Aldrich, MO, USA; Cat# T7951) were loaded alongside extracted sarkosyl insoluble samples (National Diagnostics, Hull, UK). After SDS-PAGE and immunoblotting, tau band intensity was quantified. A standard curve was generated to correlate tau concentration with band intensity, using the formula: x=y - c/m, where x is the amount of tau, y is the band intensity, c is the y-intercept, and m is the slope of the standard curve.

#### Immunohistochemistry and analysis of human brain tissue

Formalin fixed, paraffin-embedded brain sections (7 µm) from post-mortem human samples were deparaffinized, rehydrated, and antigen retrieval conducted using a sodium citrate buffer (10 mM trisodium citrate dihydrate, 0.05% Tween-20, pH 6.0). After washing in Tris-Buffered Saline (TBS), sections were blocked with 10% normal goat serum (NGS) in TBS for 1 hour at room temperature. Primary antibodies used were anti-S100B (1:500, Proteintech, Cat# 15146-1-AP), anti-GFAP (1:500, Invitrogen, Cat# PA1-10004), and anti-phosphorylated tau (AT8; 1:300, Thermo Fisher Scientific, Cat# MN1020). Slides were incubated with primary antibodies overnight at 4°C. After washing in TBS, secondary antibodies (Goat anti-Rabbit Alexa Fluor 488, Thermo Fisher Scientific, Cat# A-11034; Goat anti-Chicken Alexa Fluor 568, Thermo Fisher Scientific, Cat# A-11041; Goat anti-Mouse Alexa Fluor 680, Thermo Fisher Scientific, Cat# A-21057) were applied for 1 hour at room temperature in the dark.To reduce autofluorescence, slides were treated with Sudan Black solution, followed by washing in TBS. Nuclei were counterstained with Hoechst 33342 (Thermo Fisher Scientific, Cat# H3570) before coverslipping with ProLong™ Gold Antifade Mountant (Thermo Fisher Scientific, MA, USA). Slides were scanned using an Olympus VS200 Research Slide Scanner equipped with a high-resolution digital camera and fluorescence capabilities, using a 40X objective lens. The system was set to detect Hoechst, Alexa Fluor 488, Alexa Fluor 568, and Alexa Fluor 680 channels with optimal exposure times to avoid bleed-through and cross-talk between fluorophores. Using Visiopharm software, regions of interest were manually selected in the gray matter. A nuclei detection protocol was used to identify cells, followed by the segmentation of cell boundaries to encompass the entire cell body. Positive GFAP and S100B fluorescence were used to differentiate astrocytes from other cell types. AT8 fluorescence intensity was specifically quantified within astrocytes to assess tau uptake, along with GFAP and S100B intensities. Data were exported in Excel format for further statistical analysis using GraphPad Prism.

#### iPSC differentiation into NPCs

iPSCs were cultured and differentiated into neural progenitor cells (NPCs) using the Gibco™ PSC Neural Induction Medium system (Thermo Fisher Scientific). When iPSCs reached 15-25% confluency, the medium was replaced with Neural Induction Medium (NI medium), consisting of Neurobasal Medium and Neural Induction Supplement (1:50). Cells were maintained in NI medium at 37°C in a non-hypoxic incubator (20% O_2_, 5% CO_2_) for 7 days. Medium changes were performed as follows: on day 3, fresh NI medium was added at 2.5 mL per well; on day 5, the volume was doubled to 5 mL per well; on day 7, the medium was refreshed. By day 8, cells reached confluency and were ready for passaging. For passaging, medium was aspirated, and cells were dissociated using room temperature StemPro™ Accutase™ solution (Thermo Fisher Scientific, Cat# A1110501), transferred to a tube containing prewarmed DMEM/F-12 medium (Thermo Fisher Scientific, Cat# 11320033) and centrifuged at 190 x g for 2 minutes. The cell pellet was resuspended in Neural Expansion Medium (NE medium), a 1:1 mix of Neurobasal® Medium and Advanced™ DMEM⁄F-12 medium with Neural Induction Supplement. RevitaCell™ supplement (1:100, Thermo Fisher Scientific, Cat# A2644501) was added to NE medium to enhance cell survival during passaging. Cells were replated at a 1:3 ratio on Geltrex™-coated 6-well plates (Thermo Fisher Scientific, Cat# A1413201) and maintained in NE medium. The medium was refreshed every 48 hours. NPCs were passaged four times before being used for further differentiation or cryopreserved for future use.

#### NPC differentiation into astrocytes

NPCs were differentiated into astrocytes using Astrocyte Medium (ScienCell, Cat# 1801). NPCs were plated at a low density (15,000 cells/cm²) on Geltrex™-coated 6-well plates (Thermo Fisher Scientific, Cat# A1413201) in Neural Expansion Medium (NE medium) with added RevitaCell™ supplement (1:100, Thermo Fisher Scientific, Cat# A2644501). After 24 hours, the medium was replaced with Astrocyte Medium, which contains basal medium, fetal bovine serum (1:50), and astrocyte growth supplement (1:100). The medium was refreshed after another 24 hours and subsequently every 48 hours at 2.5 mL per well. By approximately day 6, when cells reached 90% confluency, they were passaged at the same density. Starting from day 30, fetal bovine serum was removed from the Astrocyte Medium. Cells were passaged at a 1:3 ratio approximately once a week. Astrocytes were maintained in these conditions up to day 60, at which point they were either cryopreserved or used for experimental assays.

#### Astrocyte exposure to human tau

Day 60 astrocytes were plated as single cell suspensions at low density (6000 cells/cm^2^) in Astrocyte medium (ScienCell, CA, USA) with RevitaCell™ supplement (1:100) and 5 µM AraC to reduce proliferation. After 24 hours, media was changed for Astrocyte medium with 5 µM AraC only. After a further 48 hours, media was changed to Astrocyte medium only. Sarkosyl-insoluble AD tau or equivalent control brain fractions were added to Astrocyte medium at a tau concentration of 35 ng/mL. For control samples with low tau concentration, a median equivalent of AD sample volumes was used. Media was exchanged for the same after 3 days. Astrocytes were incubated in spiked medium for 1, 3, 5 or 7 days for 7-days, and all astrocytes were fixed at the same timepoint in ice cold methanol for 5 min at -20°C. For RNA and ELISA analysis at day 7, media was collected and cells lysed in TRIzol reagent as described below. For astrocyte characterization, cells were lysed or fixed at day 70. To study tau handling beyond 7 days, wells were aspirated, washed and fresh Astrocyte medium added every 7 days. Plates were fixed after an additional 14 or 28 days.

#### Analysis of tau handling by iPSC-astrocytes

iPSC-derived astrocytes were cultured on PhenoPlate™ 96-well microplates (Perkin Elmer, MA, USA; Cat# 6055302) were fixed by replacing the medium with ice-cold methanol and placing at - 20°C for 5 minutes. Non-specific antibody binding was blocked with 5% bovine serum albumin (BSA; Cat# A9418 Sigma-Aldrich, MO, USA) in DPBS containing calcium and magnesium, for 1 hour at ambient temperature.

Primary antibodies against phosphorylated tau (AT8; 1:500, Thermo Fisher Scientific, Cat# MN1020), GFAP (1:500, Santa Cruz Biotechnology, Cat# sc-33673), and S100B (1:500, Proteintech, Cat# 15146-1-AP) were incubated along with nano secondary antibodies of alpaca anti-mouse IgG1 Alexa Fluor 488 (1:1000, Proteintech, Cat# sms1AF488-1), anti-mouse IgG2b Alexa Fluor 568 (1:1000, Proteintech, Cat# sms2bAF568-1), and anti-rabbit IgG Alexa Fluor 647 (1:1000, Proteintech, Cat# srbAF647-1) overnight at 4°C. Cells were washed and nuclei were counterstained with Hoechst 33342 (Thermo Fisher Scientific, Cat# H3570). Imaging was performed using a confocal Opera Phenix high-content screening system (Perkin Elmer, MA, USA). A 20X water objective lens (NA 1.0) was used, with laser excitation at 385 nm, 488 nm, 561 nm, and 640 nm for detecting Hoechst, Alexa Fluor 488, 568, and 647 fluorophores. Images covered up to 25 fields per well, with 15 z-stacks at 0.8 µm intervals. Quantification of tau uptake was conducted using Harmony software (Perkin Elmer, MA, USA). The ‘Find Nuclei’ function identified nuclear staining with Hoechst 33342. Nuclei were classified into ‘Dying cells’ and ‘Healthy cells’ using the ‘Linear Classifier’ method, based on intensity and morphology (Kerr et al., 1972). Astrocyte cell bodies were segmented using the ‘Find Cytoplasm’ function, identifying S100B and GFAP-positive cells. AT8 fluorescence intensity (maximum and average) was measured within GFAP-positive astrocytes to assess tau uptake. The ‘Find Region’ function was optimized to detect larger tau aggregates, enabling the calculation of volume and intensity properties for AT8, GFAP, and S100B channels. These parameters allowed for detailed analysis of tau uptake and astrocyte characteristics.

#### SDS-PAGE and Immunoblotting

Protein samples were prepared in 2X Sample Buffer or 4X NuPage Sample Buffer with reducing agent, then denatured by heating at 95°C for 5 minutes (or at 70°C for 10 minutes for NuPage samples). Proteins were separated on NuPAGE™ Bis-Tris 4-12% precast gels in the XCell SureLock™ Mini-Cell system using MOPS-SDS or MES-SDS running buffer, at a constant voltage of 120V. A protein ladder, Precision Plus Protein™ WesternC™ Blotting Standards (Bio-Rad, CA, USA), was loaded alongside the samples for molecular weight reference. Following electrophoresis, proteins were transferred onto Amersham™ Protran® 0.45 µm nitrocellulose membranes (Cytiva, Amersham, UK) using the XCell II™ Blot Module in transfer buffer (2 mM Tris-Base, 192 mM glycine, 20% methanol) at a constant 0.3 A for 1 hour. Non-specific binding was blocked with LI-COR TBS Blocking Buffer (LI-COR Biosciences, Lincoln, NE, USA) for 1 hour at room temperature. Primary antibodies used for detection were PHF1 (pSer396/404) tau (1:1000, Peter Davies, Cat# PHF1; RRID: AB_2315150) and anti-tau (Dako, 1:1000, Agilent, Cat# A0024; RRID: AB_10013724). Membranes were incubated with primary antibodies overnight at 4°C, followed by washing in TBS-T (Tris-buffered saline with 0.1% Tween-20). Secondary antibodies were IRDye® 680RD (1:10,000, LI-COR Biosciences, Cat# 926-68071) and IRDye® 800CW (1:10,000, LI-COR Biosciences, Cat# 926-32211) for detection. Membranes were incubated with secondary antibodies for 1 hour at room temperature, followed by washing in TBS-T. Blots were visualized using the Odyssey CLx Imaging System (LI-COR Biosciences), and band intensities were quantified using Image Studio Lite software (LI-COR Biosciences).

#### RNA extraction, RT and qPCR

Total RNA was extracted from astrocytes using TRIzol™ Reagent (Cat# 15596026) and Phasemaker™ Tubes (Cat# A33248), following the manufacturer’s protocol with the addition of glycogen to enhance RNA pellet recovery. Purified RNA (1-5 µg) was reverse transcribed into complementary DNA (cDNA) using the Maxima H Minus First Strand cDNA Synthesis Kit with dsDNase (Cat# K1681), following the manufacturer’s instructions without random hexamer primers. RT-qPCR was performed using PowerUp™ SYBR™ Green Master Mix on a QuantStudio™ 7 Flex Real-Time PCR System. For 96-well PCR plates, each reaction contained 10 µL of master mix, 2 µL of 5 µM primers (combined forward and reverse), 2 µL of cDNA, and 8 µL of nuclease-free water, totaling 20 µL per reaction. For 384-well plates, reaction volumes were halved to 10 µL.

Gene expression analysis was conducted using the comparative CT method, normalized to two internal control genes, β-ACTIN and GAPDH, relative to a control sample. The fluorescence threshold was set automatically using QuantStudio™ Real-Time PCR Software v1.7.1. No template controls (NTC) were included to detect contamination or primer-dimer formation, and ‘Undetermined’ Ct values in NTCs were confirmed before analyzing sample data.

#### Liquid Chromatography-Tandem Mass Spectrometry (LC-MS/MS)

Samples were submitted to the CEMS Proteomics Facility (James Black Centre, King’s College London) for analysis by high resolution Orbitrap tandem mass spectrometry coupled to liquid chromatography for protein identification. Sarkosyl-insoluble pellets from the temporal cortex of AD and control patients were sonicated in DPBS, and protein concentration was determined using a BCA assay. Samples (30 µL) were prepared for analysis by high-resolution Orbitrap tandem mass spectrometry. For enzymatic digestion, 70 µL of 50 mM triethylammonium bicarbonate (TEAB, Cat. No. T7408; Merck) was added to each sample to a total volume of 100 µL. After brief vortexing, 11 µL of 50 mM dithiothreitol (DTT, Cat. No. D5545; Merck) was added, followed by incubation at 56°C for 30 minutes. For alkylation, 12 µL of 200 mM iodoacetamide (IAA, Cat. No. I1149; Merck) was added and incubated at room temperature in the dark for 20 minutes. The reaction was quenched with 5 µL of 50 mM DTT. Finally, 1 µg of trypsin (Cat. No. 000000011047841001; Merck) was added, and samples were incubated overnight at 37°C. Peptides were dried using a Speedvac (Thermo Fisher Scientific), resuspended in 0.1% TFA, and purified using C18 spin columns (#89852; Thermo Fisher Scientific). Peptides were eluted in 50% acetonitrile/0.1% TFA, dried again, and stored at -80°C. Peptides were resuspended in MS sample buffer and injected for analysis on a U3000 UHPLC NanoLC system (Thermo Fisher Scientific, UK). Peptide separation was performed on a 75 mm C18 Pepmap column (50 cm length) using a linear gradient of 80% acetonitrile in 0.1% formic acid, with a flow rate of 250 nL/min over 60 minutes. Electrospray ionization was performed using an Orbitrap Fusion Lumos (Thermo Fisher Scientific, UK). Full MS scans (FTMS1) were acquired at a resolution of 120,000 over an m/z range of 375-1800. MS/MS fragmentation (ITMS2) used collision-induced dissociation with a 3-second cycle time, dynamic exclusion of 35 seconds, and isolation width of 1.6 m/z. The AGC target was set to 4.0e5 for FTMS1 and 1.0e4 for ITMS2, with a maximum injection time of 35 ms.

#### Analysis of LC-MS/MS data

Raw mass spectrometry data were processed into peak list files using Proteome Discoverer (ThermoScientific; v2.5) (Figure 1). The raw data file was searched using the Sequest (Eng *et al*; PMID 24226387) search algorithm against the Uniprot Human Taxonomy database (51,829 entries) and a bespoke database containing 6 tau isoforms of the human CNS (P10636-2, P10636-6, P10636-4, P10636-7, P10636-5, P10636-8). Database searching was performed at a stringency of 1% FDR including a decoy search. Posttranslational modifications for carbamidomethylation (C, static), oxidation (M, variable) and phosphorylation (S, T & Y; variable) were included in the database search. The database output file was uploaded into Scaffold software (version 5.1.2; www.proteomesoftware.com) for visualization and manual verification in the following files:

“PR710 MR3_1_3_9_HT 20230724_EDIT” and “PR710 MR3_1_3_9_Tau Isoforms 20230830_EDIT”.

The spectra of tau phosphorylation sites discovered through database searching were manually verified. To determine the relative abundance of tau peptides that were modified by phosphorylation, the ratio of modified to unmodified peptides were calculated for each phosphorylation site using their precursor ion abundance. If more than one peptide with the same phosphorylation site was detected, these precursor ion abundance values were combined and compared to unmodified peptides of the exact match. For modified peptides with no equivalent unmodified peptide and to ensure a value for each count, a small pseudo-constant at 1/10^th^ of the smallest non-zero abundance for unmodified peptides were used and applied to determine an adjusted ratio for each phosphorylation site (adjusted ratio = (modified precursor abundance)/(unmodified precursor abundance + pseudo-count). The adjusted ratio was normalized between 0 and 1 for each case and these values were plotted as a heatmap to represent the relative abundance of each phosphorylation site in each sample.

For cluster analysis, normalized values of phosphorylation sites were converted to a binary scale, where any non-zero value was designated as ’1’ (presence) and zeros were maintained as ’0’ (absence). Hierarchical clustering was performed on the binary-transformed data to explore the patterns of phosphorylation site presence across different samples. The analysis was conducted using Python’s Seaborn and Matplotlib libraries. The Ward’s method was employed for clustering, which minimizes the variance within each cluster. The Euclidean distance metric was used to quantify the dissimilarity between the data points.

Database searching of via the Uniprot Human Taxonomy database (51,829 entries) was used to determine the most abundant proteins in AD and control brain-derived sarkosyl-insoluble fractions by the precursor ion abundance. The mean of the Log2 of the precursor ion abundance was ranked for each protein and the top 20 for each sample were combined for a total of 43 proteins.

Using Python, the protein abundance data were standardized using the `StandardScaler` function from the `scikit-learn` library to normalize the values. Hierarchical clustering was performed using the Ward’s method with the `linkage` function from the `scipy` library, minimizing within-cluster variance. The clustering results were visualized through a dendrogram and a heatmap. The dendrogram was created using `scipy`, while the heatmap was generated using the `seaborn.clustermap` function, both incorporating Ward’s method and Euclidean distance. Data visualization was further enhanced using `matplotlib`.

To compare AD (n=6) and control groups (n=3), the mean of the log2 precursor ion abundance for all detected proteins and fold change between the two groups was calculated and significant differences determined by an unpaired t test. The top 20 most significantly altered proteins were plotted in a bar chart.

#### RNAseq

Cells were washed to remove cellular debris and lysed with TRIzol™ Reagent (cat# 15596026) at approximately 0.4 mL reagent per 1 x 10^5^ cells. Total RNA was extracted from lysates using Phasemaker™ Tubes (Cat# A33248), following the manufacturer’s protocol with the addition of glycogen to enhance RNA pellet recovery. The RNA concentration and purity of the resulting samples was determined using the NanoDrop™ One/OneC Microvolume UV-Vis Spectrophotometer to ensure absorbance ratio of A280/A260 of approximately 2.0 and A260/A230 ratio above 2.0. Extracted RNA was sent on dry ice to Genewiz by Azenta Life Sciences in Takely, UK for further RNA QC, library preparation (Illumina RNA with PolyA selection), sequencing (Illumina NovaSeq 2x150bp, 350M read pairs). Azenta provided a basic analysis package which included QC report, FASTQ files, data QC, trimming, mapping, differential gene expression, alternative splicing and gene ontology analysis. Tau uptake for RNAseq analysis was performed in 3 separate experiments and data averaged for analysis. Data was subsequently analyzed for comparison of differentially expressed genes (filtered by adjusted p value of <0.05) either by pooled AD and Control treated groups against untreated, or as individual cases against untreated.

Using the WGCNA package in R, a weighted gene co-expression network was constructed based on Pearson correlation coefficients, transformed into an adjacency matrix using an optimal soft thresholding power (β) to achieve a scale-free topology. Gene modules were identified by hierarchical clustering with the dynamic tree cut method. Each module’s expression profile was summarized by calculating the module eigengene (first principal component). Hub genes within significant modules were identified based on high connectivity and correlation with module eigengenes.

## Author contributions

WN, BGP-N, MJR and MLS designed the study and WN and BGP-N supervised the research. MJR planned and performed most experiments and analysed data with help from MLS, SL and CT with critical input from DS. All authors read, edited, and approved the final manuscript.

## Acknowledgements

We thank the late Professor Peter Davies for tau antibodies, Dr George Chennell of the Wohl Cellular Imaging Centre at King’s College London for technical support, and the London Neurodegenerative Disease Brain Bank.

## References

1. Ajoolabady, A., Lindholm, D., Ren, J., & Pratico, D. (2022). ER stress and UPR in Alzheimer’s disease: mechanisms, pathogenesis, treatments. Cell Death Dis, 13(8), 706. 10.1038/s41419-022-05153-5

2. Alberini, C. M. (2023). IGF2 in memory, neurodevelopmental disorders, and neurodegenerative diseases. Trends Neurosci, 46(6), 488–502. 10.1016/j.tins.2023.03.007

3. Annabi, B., Zgheib, A., & Annabi, B. (2017). Cavin-2 Functions as a Suppressive Regulator in TNF-induced Mesenchymal Stromal Cell Inflammation and Angiogenic Phenotypes. Int J Stem Cells, 10(1), 103–113. 10.15283/ijsc16032

4. Atanasova, V. S., Russell, R. J., Webster, T. G., Cao, Q., Agarwal, P., Lim, Y. Z., Krishnan, S., Fuentes, I., Guttmann-Gruber, C., McGrath, J. A., Salas-Alanis, J. C., Fertala, A., & South, A. P. (2019). Thrombospondin-1 Is a Major Activator of TGF-beta Signaling in Recessive Dystrophic Epidermolysis Bullosa Fibroblasts. J Invest Dermatol, 139(7), 1497–1505 e1495. 10.1016/j.jid.2019.01.011

5. Averna, M., De Tullio, R., Capini, P., Salamino, F., Pontremoli, S., & Melloni, E. (2003). Changes in calpastatin localization and expression during calpain activation: a new mechanism for the regulation of intracellular Ca(2+)-dependent proteolysis. Cell Mol Life Sci, 60(12), 2669–2678. 10.1007/s00018-003-3288-0

6. Bai, B., Hales, C. M., Chen, P. C., Gozal, Y., Dammer, E. B., Fritz, J. J., Wang, X., Xia, Q., Duong, D. M., Street, C., Cantero, G., Cheng, D., Jones, D. R., Wu, Z., Li, Y., Diner, I., Heilman, C. J., Rees, H. D., Wu, H., … Peng, J. (2013). U1 small nuclear ribonucleoprotein complex and RNA splicing alterations in Alzheimer’s disease. Proc Natl Acad Sci U S A, 110(41), 16562–16567. 10.1073/pnas.1310249110

7. Biondo, E. D., Spontarelli, K., Ababioh, G., Mendez, L., & Artigas, P. (2021). Diseases caused by mutations in the Na(+)/K(+) pump alpha1 gene ATP1A1. Am J Physiol Cell Physiol, 321(2), C394–C408. 10.1152/ajpcell.00059.2021

8. Bishof, I., Dammer, E. B., Duong, D. M., Kundinger, S. R., Gearing, M., Lah, J. J., Levey, A. I., & Seyfried, N. T. (2018). RNA-binding proteins with basic-acidic dipeptide (BAD) domains self-assemble and aggregate in Alzheimer’s disease. J Biol Chem, 293(28), 11047–11066. 10.1074/jbc.RA118.001747

9. Bivona, G., Iemmolo, M., & Ghersi, G. (2023). CX3CL1 Pathway as a Molecular Target for Treatment Strategies in Alzheimer’s Disease. Int J Mol Sci, 24(9). 10.3390/ijms24098230

10. Blair, L. J., Baker, J. D., Sabbagh, J. J., & Dickey, C. A. (2015). The emerging role of peptidyl-prolyl isomerase chaperones in tau oligomerization, amyloid processing, and Alzheimer’s disease. J Neurochem, 133(1), 1–13. 10.1111/jnc.13033

11. Brandebura, A. N., Paumier, A., Onur, T. S., & Allen, N. J. (2023). Astrocyte contribution to dysfunction, risk and progression in neurodegenerative disorders. Nat Rev Neurosci, 24(1), 23–39. 10.1038/s41583-022-00641-1

12. Brown, C. A., Schmidt, C., Poulter, M., Hummerich, H., Klohn, P. C., Jat, P., Mead, S., Collinge, J., & Lloyd, S. E. (2014). In vitro screen of prion disease susceptibility genes using the scrapie cell assay. Hum Mol Genet, 23(19), 5102–5108. 10.1093/hmg/ddu233

13. Burda, J. E., O’Shea, T. M., Ao, Y., Suresh, K. B., Wang, S., Bernstein, A. M., Chandra, A., Deverasetty, S., Kawaguchi, R., Kim, J. H., McCallum, S., Rogers, A., Wahane, S., & Sofroniew, M. V. (2022). Divergent transcriptional regulation of astrocyte reactivity across disorders. Nature, 606(7914), 557–564. 10.1038/s41586-022-04739-5

14. Cassano, T., Giamogante, F., Calcagnini, S., Romano, A., Lavecchia, A. M., Inglese, F., Paglia, G., Bukke, V. N., Romano, A. D., Friuli, M., Altieri, F., & Gaetani, S. (2023). PDIA3 Expression Is Altered in the Limbic Brain Regions of Triple-Transgenic Mouse Model of Alzheimer’s Disease. Int J Mol Sci, 24(3). 10.3390/ijms24033005

15. Chai, H., Diaz-Castro, B., Shigetomi, E., Monte, E., Octeau, J. C., Yu, X., Cohn, W., Rajendran, P. S., Vondriska, T. M., Whitelegge, J. P., Coppola, G., & Khakh, B. S. (2017). Neural Circuit-Specialized Astrocytes: Transcriptomic, Proteomic, Morphological, and Functional Evidence. Neuron, 95(3), 531–549 e539. 10.1016/j.neuron.2017.06.029

16. Chen, D., Drombosky, K. W., Hou, Z., Sari, L., Kashmer, O. M., Ryder, B. D., Perez, V. A., Woodard, D. R., Lin, M. M., Diamond, M. I., & Joachimiak, L. A. (2019). Tau local structure shields an amyloid-forming motif and controls aggregation propensity. Nat Commun, 10(1), 2493. 10.1038/s41467-019-10355-1

17. Chiariello, A., Valente, S., Pasquinelli, G., Baracca, A., Sgarbi, G., Solaini, G., Medici, V., Fantini, V., Poloni, T. E., Tognocchi, M., Arcaro, M., Galimberti, D., Franceschi, C., Capri, M., Salvioli, S., & Conte, M. (2022). The expression pattern of GDF15 in human brain changes during aging and in Alzheimer’s disease. Front Aging Neurosci, 14, 1058665. 10.3389/fnagi.2022.1058665

18. Choi, H., Lee, E. J., Shin, J. S., Kim, H., Bae, S., Choi, Y., & Lee, D. S. (2023). Spatiotemporal characterization of glial cell activation in an Alzheimer’s disease model by spatially resolved transcriptomics. Exp Mol Med. 10.1038/s12276-023-01123-9

19. Cicognola, C., Satir, T. M., Brinkmalm, G., Matecko-Burmann, I., Agholme, L., Bergstrom, P., Becker, B., Zetterberg, H., Blennow, K., & Hoglund, K. (2020). Tauopathy-Associated Tau Fragment Ending at Amino Acid 224 Is Generated by Calpain-2 Cleavage. J Alzheimers Dis, 74(4), 1143–1156. 10.3233/JAD-191130

20. Cocks, G., Curran, S., Gami, P., Uwanogho, D., Jeffries, A. R., Kathuria, A., Lucchesi, W., Wood, V., Dixon, R., Ogilvie, C., Steckler, T., & Price, J. (2013). The utility of patient specific induced pluripotent stem cells for the modelling of Autistic Spectrum Disorders. Psychopharmacology, 231(6), 1079–1088. 10.1007/s00213-013-3196-4

21. Crowther, R. A., & Goedert, M. (2000). Abnormal tau-containing filaments in neurodegenerative diseases. J Struct Biol, 130(2-3), 271–279. 10.1006/jsbi.2000.4270

22. Dang, Y., He, Q., Yang, S., Sun, H., Liu, Y., Li, W., Tang, Y., Zheng, Y., & Wu, T. (2022). FTH1-and SAT1-Induced Astrocytic Ferroptosis Is Involved in Alzheimer’s Disease: Evidence from Single-Cell Transcriptomic Analysis. Pharmaceuticals (Basel), 15(10). 10.3390/ph15101177

23. de Calignon, A., Polydoro, M., Suarez-Calvet, M., William, C., Adamowicz, D. H., Kopeikina, K. J., Pitstick, R., Sahara, N., Ashe, K. H., Carlson, G. A., Spires-Jones, T. L., & Hyman, B. T. (2012). Propagation of tau pathology in a model of early Alzheimer’s disease. Neuron, 73(4), 685–697. 10.1016/j.neuron.2011.11.033

24. Dujardin, S., Begard, S., Caillierez, R., Lachaud, C., Carrier, S., Lieger, S., Gonzalez, J. A., Deramecourt, V., Deglon, N., Maurage, C. A., Frosch, M. P., Hyman, B. T., Colin, M., & Buee, L. (2018). Different tau species lead to heterogeneous tau pathology propagation and misfolding. Acta Neuropathol Commun, 6(1), 132. 10.1186/s40478-018-0637-7

25. Dujardin, S., Commins, C., Lathuiliere, A., Beerepoot, P., Fernandes, A. R., Kamath, T. V., De Los Santos, M. B., Klickstein, N., Corjuc, D. L., Corjuc, B. T., Dooley, P. M., Viode, A., Oakley, D. H., Moore, B. D., Mullin, K., Jean-Gilles, D., Clark, R., Atchison, K., Moore, R., … Hyman, B. T. (2020). Tau molecular diversity contributes to clinical heterogeneity in Alzheimer’s disease. Nat Med, 26(8), 1256–1263. 10.1038/s41591-020-0938-9

26. Eisenbaum, M., Pearson, A., Ortiz, C., Mullan, M., Crawford, F., Ojo, J., & Bachmeier, C. (2023). ApoE4 expression disrupts tau uptake, trafficking, and clearance in astrocytes. Glia. 10.1002/glia.24469

27. Eltom, K., Mothes, T., Libard, S., Ingelsson, M., & Erlandsson, A. (2024). Astrocytic accumulation of tau fibrils isolated from Alzheimer’s disease brains induces inflammation, cell-to-cell propagation and neuronal impairment. Acta Neuropathol Commun, 12(1), 34. 10.1186/s40478-024-01745-8

28. Escartin, C., Galea, E., Lakatos, A., O’Callaghan, J. P., Petzold, G. C., Serrano-Pozo, A., Steinhauser, C., Volterra, A., Carmignoto, G., Agarwal, A., Allen, N. J., Araque, A., Barbeito, L., Barzilai, A., Bergles, D. E., Bonvento, G., Butt, A. M., Chen, W. T., Cohen-Salmon, M., … Verkhratsky, A. (2021). Reactive astrocyte nomenclature, definitions, and future directions. Nat Neurosci, 24(3), 312–325. 10.1038/s41593-020-00783-4

29. Esposito, G., Scuderi, C., Lu, J., Savani, C., De Filippis, D., Iuvone, T., Steardo, L., Jr., Sheen, V., & Steardo, L. (2008). S100B induces tau protein hyperphosphorylation via Dickopff-1 up-regulation and disrupts the Wnt pathway in human neural stem cells. J Cell Mol Med, 12(3), 914–927. 10.1111/j.1582-4934.2008.00159.x

30. Fan, G., Li, Y., Zong, Y., Suo, X., Jia, Y., Gao, M., & Yang, X. (2023). GPAT3 regulates the synthesis of lipid intermediate LPA and exacerbates Kupffer cell inflammation mediated by the ERK signaling pathway. Cell Death Dis, 14(3), 208. 10.1038/s41419-023-05741-z

31. Fernandez-Hernando, C., & Suarez, Y. (2020). ANGPTL4: a multifunctional protein involved in metabolism and vascular homeostasis. Curr Opin Hematol, 27(3), 206–213. 10.1097/MOH.0000000000000580

32. Ferrari-Souza, J. P., Ferreira, P. C. L., Bellaver, B., Tissot, C., Wang, Y. T., Leffa, D. T., Brum, W. S., Benedet, A. L., Ashton, N. J., De Bastiani, M. A., Rocha, A., Therriault, J., Lussier, F. Z., Chamoun, M., Servaes, S., Bezgin, G., Kang, M. S., Stevenson, J., Rahmouni, N., … Pascoal, T. A. (2022). Astrocyte biomarker signatures of amyloid-beta and tau pathologies in Alzheimer’s disease. Mol Psychiatry, 27(11), 4781–4789. 10.1038/s41380-022-01716-2

33. Fitzgerald, G. S., Chuchta, T. G., & McNay, E. C. (2023). Insulin-like growth factor-2 is a promising candidate for the treatment and prevention of Alzheimer’s disease. CNS Neurosci Ther, 29(6), 1449–1469. 10.1111/cns.14160

34. Fitzpatrick, A. W. P., Falcon, B., He, S., Murzin, A. G., Murshudov, G., Garringer, H. J., Crowther, R. A., Ghetti, B., Goedert, M., & Scheres, S. H. W. (2017). Cryo-EM structures of tau filaments from Alzheimer’s disease. Nature, 547(7662), 185–190. 10.1038/nature23002

35. Frederic, M. Y., Monino, C., Marschall, C., Hamroun, D., Faivre, L., Jondeau, G., Klein, H. G., Neumann, L., Gautier, E., Binquet, C., Maslen, C., Godfrey, M., Gupta, P., Milewicz, D., Boileau, C., Claustres, M., Beroud, C., & Collod-Beroud, G. (2009). The FBN2 gene: new mutations, locus-specific database (Universal Mutation Database FBN2), and genotype-phenotype correlations. Hum Mutat, 30(2), 181–190. 10.1002/humu.20794

36. Gentil, B. J., Delphin, C., Mbele, G. O., Deloulme, J. C., Ferro, M., Garin, J., & Baudier, J. (2001). The giant protein AHNAK is a specific target for the calcium- and zinc-binding S100B protein: potential implications for Ca2+ homeostasis regulation by S100B. J Biol Chem, 276(26), 23253–23261. 10.1074/jbc.M010655200

37. Ghafouri-Fard, S., Khoshbakht, T., Taheri, M., & Ghanbari, M. (2021). A concise review on the role of BDNF-AS in human disorders. Biomed Pharmacother, 142, 112051. 10.1016/j.biopha.2021.112051

38. Ghasempour, G., Mohammadi, A., Zamani-Garmsiri, F., Soleimani, A. A., & Najafi, M. (2022). Upregulation of TGF-beta type II receptor in high glucose-induced vascular smooth muscle cells. Mol Biol Rep, 49(4), 2869–2875. 10.1007/s11033-021-07100-7

39. Gns, H. S., Rajalekshmi, S. G., & Burri, R. R. (2022). Revelation of Pivotal Genes Pertinent to Alzheimer’s Pathogenesis: A Methodical Evaluation of 32 GEO Datasets. J Mol Neurosci, 72(2), 303–322. 10.1007/s12031-021-01919-2

40. Goedert, M., Eisenberg, D. S., & Crowther, R. A. (2017). Propagation of Tau Aggregates and Neurodegeneration. Annu Rev Neurosci, 40, 189–210. 10.1146/annurev-neuro-072116-031153

41. Gregorio, I., Braghetta, P., Bonaldo, P., & Cescon, M. (2018). Collagen VI in healthy and diseased nervous system. Dis Model Mech, 11(6). 10.1242/dmm.032946

42. Habib, N., McCabe, C., Medina, S., Varshavsky, M., Kitsberg, D., Dvir-Szternfeld, R., Green, G., Dionne, D., Nguyen, L., Marshall, J. L., Chen, F., Zhang, F., Kaplan, T., Regev, A., & Schwartz, M. (2020). Disease-associated astrocytes in Alzheimer’s disease and aging. Nat Neurosci, 23(6), 701–706. 10.1038/s41593-020-0624-8

43. Haj-Yahya, M., Gopinath, P., Rajasekhar, K., Mirbaha, H., Diamond, M. I., & Lashuel, H. A. (2020). Site-Specific Hyperphosphorylation Inhibits, Rather than Promotes, Tau Fibrillization, Seeding Capacity, and Its Microtubule Binding. Angew Chem Int Ed Engl, 59(10), 4059–4067. 10.1002/anie.201913001

44. Han, C., Yan, P., He, T., Cheng, J., Zheng, W., Zheng, L. T., & Zhen, X. (2020). PHLDA1 promotes microglia-mediated neuroinflammation via regulating K63-linked ubiquitination of TRAF6. Brain Behav Immun, 88, 640–653. 10.1016/j.bbi.2020.04.064

45. He, Y., Kaya, I., Shariatgorji, R., Lundkvist, J., Wahlberg, L. U., Nilsson, A., Mamula, D., Kehr, J., Zareba-Paslawska, J., Biverstal, H., Chergui, K., Zhang, X., Andren, P. E., & Svenningsson, P. (2023). Prosaposin maintains lipid homeostasis in dopamine neurons and counteracts experimental parkinsonism in rodents. Nat Commun, 14(1), 5804. 10.1038/s41467-023-41539-5

46. Hu, D., Sun, X., Magpusao, A., Fedorov, Y., Thompson, M., Wang, B., Lundberg, K., Adams, D. J., & Qi, X. (2021). Small-molecule suppression of calpastatin degradation reduces neuropathology in models of Huntington’s disease. Nat Commun, 12(1), 5305. 10.1038/s41467-021-25651-y

47. Hu, L. T., Xie, X. Y., Zhou, G. F., Wen, Q. X., Song, L., Luo, B., Deng, X. J., Pan, Q. L., & Chen, G. J. (2023). HMGCS2-Induced Autophagic Degradation of Tau Involves Ketone Body and ANKRD24. J Alzheimers Dis, 91(1), 407–426. 10.3233/JAD-220640

48. Hu, L. T., Zhu, B. L., Lai, Y. J., Long, Y., Zha, J. S., Hu, X. T., Zhang, J. H., & Chen, G. J. (2017). HMGCS2 promotes autophagic degradation of the amyloid-beta precursor protein through ketone body-mediated mechanisms. Biochem Biophys Res Commun, 486(2), 492–498. 10.1016/j.bbrc.2017.03.069

49. Huang, X., Liu, L., Yao, J., Lin, C., Xiang, T., & Yang, A. (2023). S-acylation regulates SQSTM1/p62-mediated selective autophagy. Autophagy, 1-3. 10.1080/15548627.2023.2297623

50. Jasmin, J. F., Frank, P. G., & Lisanti, M. P. (2012). CAVEOLIN-1: Role in Cell Signaling. In Caveolins and Caveolae (Vol. 723, pp. 29–50). Springer. 10.1007/978-1-4614-1222-9_3

51. Jensen, H. L. B., Lillenes, M. S., Rabano, A., Gunther, C. C., Riaz, T., Kalayou, S. T., Ulstein, I. D., Bohmer, T., & Tonjum, T. (2018). Expression of nucleotide excision repair in Alzheimer’s disease is higher in brain tissue than in blood. Neurosci Lett, 672, 53–58. 10.1016/j.neulet.2018.02.043

52. Jin, X., Nie, E., Zhou, X., Zeng, A., Yu, T., Zhi, T., Jiang, K., Wang, Y., Zhang, J., & You, Y. (2017). Fstl1 Promotes Glioma Growth Through the BMP4/Smad1/5/8 Signaling Pathway. Cell Physiol Biochem, 44(4), 1616–1628. 10.1159/000485759

53. Jiwaji, Z., Tiwari, S. S., Aviles-Reyes, R. X., Hooley, M., Hampton, D., Torvell, M., Johnson, D. A., McQueen, J., Baxter, P., Sabari-Sankar, K., Qiu, J., He, X., Fowler, J., Febery, J., Gregory, J., Rose, J., Tulloch, J., Loan, J., Story, D., … Hardingham, G. E. (2022). Reactive astrocytes acquire neuroprotective as well as deleterious signatures in response to Tau and Ass pathology. Nat Commun, 13(1), 135. 10.1038/s41467-021-27702-w

54. Kageyama, S., Gudmundsson, S. R., Sou, Y. S., Ichimura, Y., Tamura, N., Kazuno, S., Ueno, T., Miura, Y., Noshiro, D., Abe, M., Mizushima, T., Miura, N., Okuda, S., Motohashi, H., Lee, J. A., Sakimura, K., Ohe, T., Noda, N. N., Waguri, S., … Komatsu, M. (2021). p62/SQSTM1-droplet serves as a platform for autophagosome formation and anti-oxidative stress response. Nat Commun, 12(1), 16. 10.1038/s41467-020-20185-1

55. Kamath, T. V., Klickstein, N., Commins, C., Fernandes, A. R., Oakley, D. H., Frosch, M. P., Hyman, B. T., & Dujardin, S. (2021). Kinetics of tau aggregation reveals patient-specific tau characteristics among Alzheimer’s cases. Brain Commun, 3(2), fcab096. 10.1093/braincomms/fcab096

56. Karch, C. M., Ezerskiy, L. A., Bertelsen, S., Alzheimer’s Disease Genetics, C., & Goate, A. M. (2016). Alzheimer’s Disease Risk Polymorphisms Regulate Gene Expression in the ZCWPW1 and the CELF1 Loci. PLoS One, 11(2), e0148717. 10.1371/journal.pone.0148717

57. Karlsson, M., Zhang, C., Mear, L., Zhong, W., Digre, A., Katona, B., Sjostedt, E., Butler, L., Odeberg, J., Dusart, P., Edfors, F., Oksvold, P., von Feilitzen, K., Zwahlen, M., Arif, M., Altay, O., Li, X., Ozcan, M., Mardinoglu, A., … Lindskog, C. (2021). A single-cell type transcriptomics map of human tissues. Sci Adv, 7(31). 10.1126/sciadv.abh2169

58. Kerr, J. F., Wyllie, A. H., & Currie, A. R. (1972). Apoptosis: a basic biological phenomenon with wide-ranging implications in tissue kinetics. Br J Cancer, 26(4), 239–257. 10.1038/bjc.1972.33

59. Kini, H. K., Kong, J., & Liebhaber, S. A. (2014). Cytoplasmic poly(A) binding protein C4 serves a critical role in erythroid differentiation. Mol Cell Biol, 34(7), 1300–1309. 10.1128/MCB.01683-13

60. Konishi, H., Koizumi, S., & Kiyama, H. (2022). Phagocytic astrocytes: Emerging from the shadows of microglia. Glia, 70(6), 1009–1026. 10.1002/glia.24145

61. Kovacs, G. G. (2020). Astroglia and Tau: New Perspectives. Front Aging Neurosci, 12, 96. 10.3389/fnagi.2020.00096

62. Kulczynska-Przybik, A., Mroczko, P., Dulewicz, M., & Mroczko, B. (2021). The Implication of Reticulons (RTNs) in Neurodegenerative Diseases: From Molecular Mechanisms to Potential Diagnostic and Therapeutic Approaches. Int J Mol Sci, 22(9). 10.3390/ijms22094630

63. Kurbatskaya, K., Phillips, E. C., Croft, C. L., Dentoni, G., Hughes, M. M., Wade, M. A., Al-Sarraj, S., Troakes, C., O’Neill, M. J., Perez-Nievas, B. G., Hanger, D. P., & Noble, W. (2016). Upregulation of calpain activity precedes tau phosphorylation and loss of synaptic proteins in Alzheimer’s disease brain. Acta Neuropathol Commun, 4, 34. 10.1186/s40478-016-0299-2

64. Lau, L. W., Cua, R., Keough, M. B., Haylock-Jacobs, S., & Yong, V. W. (2013). Pathophysiology of the brain extracellular matrix: a new target for remyelination. Nat Rev Neurosci, 14(10), 722–729. 10.1038/nrn3550

65. Liu, Z., Lai, J., Jiang, H., Ma, C., & Huang, H. (2021). Collagen XI alpha 1 chain, a potential therapeutic target for cancer. FASEB J, 35(6), e21603. 10.1096/fj.202100054RR

66. Loov, C., Mitchell, C. H., Simonsson, M., & Erlandsson, A. (2015). Slow degradation in phagocytic astrocytes can be enhanced by lysosomal acidification. Glia, 63(11), 1997–2009. 10.1002/glia.22873

67. Mandal, P., Belapurkar, V., Nair, D., & Ramanan, N. (2021). Vinculin-mediated axon growth requires interaction with actin but not talin in mouse neocortical neurons. Cell Mol Life Sci, 78(15), 5807–5826. 10.1007/s00018-021-03879-7

68. Manning, G., Whyte, D. B., Martinez, R., Hunter, T., & Sudarsanam, S. (2002). The protein kinase complement of the human genome. Science, 298(5600), 1912–1934. 10.1126/science.1075762

69. Martini-Stoica, H., Cole, A. L., Swartzlander, D. B., Chen, F., Wan, Y. W., Bajaj, L., Bader, D. A., Lee, V. M. Y., Trojanowski, J. Q., Liu, Z., Sardiello, M., & Zheng, H. (2018). TFEB enhances astroglial uptake of extracellular tau species and reduces tau spreading. J Exp Med, 215(9), 2355–2377. 10.1084/jem.20172158

70. Mathias, R. A., Pani, V., & Chilton, F. H. (2014). Genetic Variants in the FADS Gene: Implications for Dietary Recommendations for Fatty Acid Intake. Curr Nutr Rep, 3(2), 139–148. 10.1007/s13668-014-0079-1

71. Middeldorp, J., & Hol, E. M. (2011). GFAP in health and disease. Prog Neurobiol, 93(3), 421–443. 10.1016/j.pneurobio.2011.01.005

72. Miller, S. J. (2018). Astrocyte Heterogeneity in the Adult Central Nervous System. Front Cell Neurosci, 12, 401. 10.3389/fncel.2018.00401

73. Moreira, G. G., Cantrelle, F. X., Quezada, A., Carvalho, F. S., Cristovao, J. S., Sengupta, U., Puangmalai, N., Carapeto, A. P., Rodrigues, M. S., Cardoso, I., Fritz, G., Herrera, F., Kayed, R., Landrieu, I., & Gomes, C. M. (2021). Dynamic interactions and Ca(2+)-binding modulate the holdase-type chaperone activity of S100B preventing tau aggregation and seeding. Nat Commun, 12(1), 6292. 10.1038/s41467-021-26584-2

74. Moreira, G. G., & Gomes, C. M. (2023). Tau liquid-liquid phase separation is modulated by the Ca(2+) -switched chaperone activity of the S100B protein. J Neurochem, 166(1), 76–86. 10.1111/jnc.15756

75. Moretto, E., Stuart, S., Surana, S., Vargas, J. N. S., & Schiavo, G. (2022). The Role of Extracellular Matrix Components in the Spreading of Pathological Protein Aggregates. Front Cell Neurosci, 16, 844211. 10.3389/fncel.2022.844211

76. Mothes, T., Portal, B., Konstantinidis, E., Eltom, K., Libard, S., Streubel-Gallasch, L., Ingelsson, M., Rostami, J., Lindskog, M., & Erlandsson, A. (2023). Astrocytic uptake of neuronal corpses promotes cell-to-cell spreading of tau pathology. Acta Neuropathol Commun, 11(1), 97. 10.1186/s40478-023-01589-8

77. Nolan, A., De Paula Franca Resende, E., Petersen, C., Neylan, K., Spina, S., Huang, E., Seeley, W., Miller, Z., & Grinberg, L. T. (2019). Astrocytic Tau Deposition Is Frequent in Typical and Atypical Alzheimer Disease Presentations. J Neuropathol Exp Neurol, 78(12), 1112–1123. 10.1093/jnen/nlz094

78. Ogbu, S. C., Musich, P. R., Zhang, J., Yao, Z. Q., Howe, P. H., & Jiang, Y. (2021). The role of disabled-2 (Dab2) in diseases. Gene, 769, 145202. 10.1016/j.gene.2020.145202

79. Ono, Y., Saido, T. C., & Sorimachi, H. (2016). Calpain research for drug discovery: challenges and potential. Nat Rev Drug Discov, 15(12), 854–876. 10.1038/nrd.2016.212

80. Parast, M. M., & Otey, C. A. (2000). Characterization of palladin, a novel protein localized to stress fibers and cell adhesions. J Cell Biol, 150(3), 643–656. 10.1083/jcb.150.3.643

81. Perea, J. R., Lopez, E., Diez-Ballesteros, J. C., Avila, J., Hernandez, F., & Bolos, M. (2019). Extracellular Monomeric Tau Is Internalized by Astrocytes. Front Neurosci, 13, 442. 10.3389/fnins.2019.00442

82. Perez-Nievas, B. G., & Serrano-Pozo, A. (2018). Deciphering the Astrocyte Reaction in Alzheimer’s Disease. Front Aging Neurosci, 10, 114. 10.3389/fnagi.2018.00114

83. Probst, A., Ulrich, J., & Heitz, P. U. (1982). Senile dementia of Alzheimer type: astroglial reaction to extracellular neurofibrillary tangles in the hippocampus. An immunocytochemical and electron-microscopic study. Acta Neuropathol, 57(1), 75–79. 10.1007/BF00688880

84. Puchalska, P., & Crawford, P. A. (2017). Multi-dimensional Roles of Ketone Bodies in Fuel Metabolism, Signaling, and Therapeutics. Cell Metab, 25(2), 262–284. 10.1016/j.cmet.2016.12.022

85. Qi, Y., Wang, M., & Jiang, Q. (2022). PABPC1--mRNA stability, protein translation and tumorigenesis. Front Oncol, 12, 1025291. 10.3389/fonc.2022.1025291

86. Rahman, M. M., & Lendel, C. (2021). Extracellular protein components of amyloid plaques and their roles in Alzheimer’s disease pathology. Mol Neurodegener, 16(1), 59. 10.1186/s13024-021-00465-0

87. Reid, M. J., Beltran-Lobo, P., Johnson, L., Perez-Nievas, B. G., & Noble, W. (2020). Astrocytes in Tauopathies. Front Neurol, 11, 572850. 10.3389/fneur.2020.572850

88. Richetin, K., Steullet, P., Pachoud, M., Perbet, R., Parietti, E., Maheswaran, M., Eddarkaoui, S., Begard, S., Pythoud, C., Rey, M., Caillierez, R., K, Q. D., Halliez, S., Bezzi, P., Buee, L., Leuba, G., Colin, M., Toni, N., & Deglon, N. (2020). Tau accumulation in astrocytes of the dentate gyrus induces neuronal dysfunction and memory deficits in Alzheimer’s disease. Nat Neurosci, 23(12), 1567–1579. 10.1038/s41593-020-00728-x

89. Robbins, J. P., Perfect, L., Ribe, E. M., Maresca, M., Dangla-Valls, A., Foster, E. M., Killick, R., Nowosiad, P., Reid, M. J., Polit, L. D., Nevado, A. J., Ebner, D., Bohlooly, Y. M., Buckley, N., Pangalos, M. N., Price, J., & Lovestone, S. (2018). Clusterin Is Required for beta-Amyloid Toxicity in Human iPSC-Derived Neurons. Front Neurosci, 12, 504. 10.3389/fnins.2018.00504

90. Sadick, J. S., O’Dea, M. R., Hasel, P., Dykstra, T., Faustin, A., & Liddelow, S. A. (2022). Astrocytes and oligodendrocytes undergo subtype-specific transcriptional changes in Alzheimer’s disease. Neuron, 110(11), 1788–1805 e1710. 10.1016/j.neuron.2022.03.008

91. Sanders, D. W., Kaufman, S. K., DeVos, S. L., Sharma, A. M., Mirbaha, H., Li, A., Barker, S. J., Foley, A. C., Thorpe, J. R., Serpell, L. C., Miller, T. M., Grinberg, L. T., Seeley, W. W., & Diamond, M. I. (2014). Distinct tau prion strains propagate in cells and mice and define different tauopathies. Neuron, 82(6), 1271–1288. 10.1016/j.neuron.2014.04.047

92. Santello, M., Toni, N., & Volterra, A. (2019). Astrocyte function from information processing to cognition and cognitive impairment. Nat Neurosci, 22(2), 154–166. 10.1038/s41593-018-0325-8

93. Sawada, R., Nakano-Doi, A., Matsuyama, T., Nakagomi, N., & Nakagomi, T. (2020). CD44 expression in stem cells and niche microglia/macrophages following ischemic stroke. Stem Cell Investig, 7, 4. 10.21037/sci.2020.02.02

94. Serrano-Pozo, A., Li, Z., Woodbury, M. E., Muñoz-Castro, C., Wachter, A., Jayakumar, R., Bryant, A. G., Noori, A., Welikovitch, L. A., Hu, M., Zhao, K., Liao, F., Lin, G., Pastika, T., Tamm, J., Abdourahman, A., Kwon, T., Bennett, R. E., Talanian, R. V., … Das, S. (2022). Astrocyte transcriptomic changes along the spatiotemporal progression of Alzheimer’s disease. BioRxiv. 10.1101/2022.12.03.518999

95. Serrano-Pozo, A., Mielke, M. L., Gomez-Isla, T., Betensky, R. A., Growdon, J. H., Frosch, M. P., & Hyman, B. T. (2011). Reactive glia not only associates with plaques but also parallels tangles in Alzheimer’s disease. Am J Pathol, 179(3), 1373–1384. 10.1016/j.ajpath.2011.05.047

96. Seto-Salvia, N., Esteras, N., de Silva, R., de Pablo-Fernandez, E., Arber, C., Toomey, C. E., Polke, J. M., Morris, H. R., Rohrer, J. D., Abramov, A. Y., Patani, R., Wray, S., & Warner, T. T. (2022). Elevated 4R-tau in astrocytes from asymptomatic carriers of the MAPT 10+16 intronic mutation. J Cell Mol Med, 26(4), 1327–1331. 10.1111/jcmm.17136

97. Sheng, J. G., Mrak, R. E., & Griffin, W. S. (1994). S100 beta protein expression in Alzheimer disease: potential role in the pathogenesis of neuritic plaques. J Neurosci Res, 39(4), 398–404. 10.1002/jnr.490390406

98. Shi, Y., Zhang, W., Yang, Y., Murzin, A. G., Falcon, B., Kotecha, A., van Beers, M., Tarutani, A., Kametani, F., Garringer, H. J., Vidal, R., Hallinan, G. I., Lashley, T., Saito, Y., Murayama, S., Yoshida, M., Tanaka, H., Kakita, A., Ikeuchi, T., … Scheres, S. H. W. (2021). Structure-based classification of tauopathies. Nature, 598(7880), 359–363. 10.1038/s41586-021-03911-7

99. Simic, G., Babic Leko, M., Wray, S., Harrington, C., Delalle, I., Jovanov-Milosevic, N., Bazadona, D., Buee, L., de Silva, R., Di Giovanni, G., Wischik, C., & Hof, P. R. (2016). Tau Protein Hyperphosphorylation and Aggregation in Alzheimer’s Disease and Other Tauopathies, and Possible Neuroprotective Strategies. Biomolecules, 6(1), 6. 10.3390/biom6010006

100. Soles, A., Selimovic, A., Sbrocco, K., Ghannoum, F., Hamel, K., Moncada, E. L., Gilliat, S., & Cvetanovic, M. (2023). Extracellular Matrix Regulation in Physiology and in Brain Disease. Int J Mol Sci, 24(8). 10.3390/ijms24087049

101. Subbarayan, M. S., Joly-Amado, A., Bickford, P. C., & Nash, K. R. (2022). CX3CL1/CX3CR1 signaling targets for the treatment of neurodegenerative diseases. Pharmacol Ther, 231, 107989. 10.1016/j.pharmthera.2021.107989

102. Tcw, J., Wang, M., Pimenova, A. A., Bowles, K. R., Hartley, B. J., Lacin, E., Machlovi, S. I., Abdelaal, R., Karch, C. M., Phatnani, H., Slesinger, P. A., Zhang, B., Goate, A. M., & Brennand, K. J. (2017). An Efficient Platform for Astrocyte Differentiation from Human Induced Pluripotent Stem Cells. Stem Cell Reports, 9(2), 600–614. 10.1016/j.stemcr.2017.06.018

103. Tijms, B. M., Vromen, E. M., Mjaavatten, O., Holstege, H., Reus, L. M., van der Lee, S., Wesenhagen, K. E. J., Lorenzini, L., Vermunt, L., Venkatraghavan, V., Tesi, N., Tomassen, J., den Braber, A., Goossens, J., Vanmechelen, E., Barkhof, F., Pijnenburg, Y. A. L., van der Flier, W. M., Teunissen, C. E., … Visser, P. J. (2024). Cerebrospinal fluid proteomics in patients with Alzheimer’s disease reveals five molecular subtypes with distinct genetic risk profiles. Nat Aging. 10.1038/s43587-023-00550-7

104. Van Eldik, L. J., & Griffin, W. S. (1994). S100 beta expression in Alzheimer’s disease: relation to neuropathology in brain regions. Biochim Biophys Acta, 1223(3), 398–403. 10.1016/0167-4889(94)90101-5

105. Verbeek, M. M., Otte-Holler, I., van den Born, J., van den Heuvel, L. P., David, G., Wesseling, P., & de Waal, R. M. (1999). Agrin is a major heparan sulfate proteoglycan accumulating in Alzheimer’s disease brain. Am J Pathol, 155(6), 2115–2125. 10.1016/S0002-9440(10)65529-0

106. Voigt, S., Jungnickel, B., Hartmann, E., & Rapoport, T. A. (1996). Signal sequence-dependent function of the TRAM protein during early phases of protein transport across the endoplasmic reticulum membrane. J Cell Biol, 134(1), 25–35. 10.1083/jcb.134.1.25

107. Wegmann, S., Nicholls, S., Takeda, S., Fan, Z., & Hyman, B. T. (2016). Formation, release, and internalization of stable tau oligomers in cells. J Neurochem, 139(6), 1163–1174. 10.1111/jnc.13866

108. Wesseling, H., Mair, W., Kumar, M., Schlaffner, C. N., Tang, S., Beerepoot, P., Fatou, B., Guise, A. J., Cheng, L., Takeda, S., Muntel, J., Rotunno, M. S., Dujardin, S., Davies, P., Kosik, K. S., Miller, B. L., Berretta, S., Hedreen, J. C., Grinberg, L. T., … Steen, J. A. (2020). Tau PTM Profiles Identify Patient Heterogeneity and Stages of Alzheimer’s Disease. Cell, 183(6), 1699–1713 e1613. 10.1016/j.cell.2020.10.029

109. Winkler, J., Abisoye-Ogunniyan, A., Metcalf, K. J., & Werb, Z. (2020). Concepts of extracellular matrix remodelling in tumour progression and metastasis. Nat Commun, 11(1), 5120. 10.1038/s41467-020-18794-x

110. Xiao, D., Su, X., Gao, H., Li, X., & Qu, Y. (2021). The Roles of Lpar1 in Central Nervous System Disorders and Diseases. Front Neurosci, 15, 710473. 10.3389/fnins.2021.710473

111. Yu, W., Yu, W., Yang, Y., & Lu, Y. (2021). Exploring the Key Genes and Identification of Potential Diagnosis Biomarkers in Alzheimer’s Disease Using Bioinformatics Analysis. Front Aging Neurosci, 13, 602781. 10.3389/fnagi.2021.602781

112. Zhang, L., Wang, B., Li, L., Qian, D. M., Yu, H., Xue, M. L., Hu, M., & Song, X. X. (2017). Antiviral effects of IFIT1 in human cytomegalovirus-infected fetal astrocytes. J Med Virol, 89(4), 672–684. 10.1002/jmv.24674

113. Zhang, Y., Sloan, S. A., Clarke, L. E., Caneda, C., Plaza, C. A., Blumenthal, P. D., Vogel, H., Steinberg, G. K., Edwards, M. S., Li, G., Duncan, J. A., 3rd, Cheshier, S. H., Shuer, L. M., Chang, E. F., Grant, G. A., Gephart, M. G., & Barres, B. A. (2016). Purification and Characterization of Progenitor and Mature Human Astrocytes Reveals Transcriptional and Functional Differences with Mouse. Neuron, 89(1), 37–53. 10.1016/j.neuron.2015.11.013

114. Zhang, Y. W., Thompson, R., Zhang, H., & Xu, H. (2011). APP processing in Alzheimer’s disease. Mol Brain, 4, 3. 10.1186/1756-6606-4-3

115. Zhong, Q., Congdon, E. E., Nagaraja, H. N., & Kuret, J. (2012). Tau isoform composition influences rate and extent of filament formation. J Biol Chem, 287(24), 20711–20719. 10.1074/jbc.M112.364067

116. Zhu, Y., Zhang, M., Kelly, A. R., & Cheng, A. (2014). The carbohydrate-binding domain of overexpressed STBD1 is important for its stability and protein-protein interactions. Biosci Rep, 34(4). 10.1042/BSR20140053

117. Zinnall, U., Milek, M., Minia, I., Vieira-Vieira, C. H., Muller, S., Mastrobuoni, G., Hazapis, O. G., Del Giudice, S., Schwefel, D., Bley, N., Voigt, F., Chao, J. A., Kempa, S., Huttelmaier, S., Selbach, M., & Landthaler, M. (2022). HDLBP binds ER-targeted mRNAs by multivalent interactions to promote protein synthesis of transmembrane and secreted proteins. Nat Commun, 13(1), 2727. 10.1038/s41467-022-30322-7

