## Supplementary figures and images for "Properties of Alzheimer’s disease brain-derived tau aggregates define tau processing by human astrocytes"

### Supplementary Fig 1

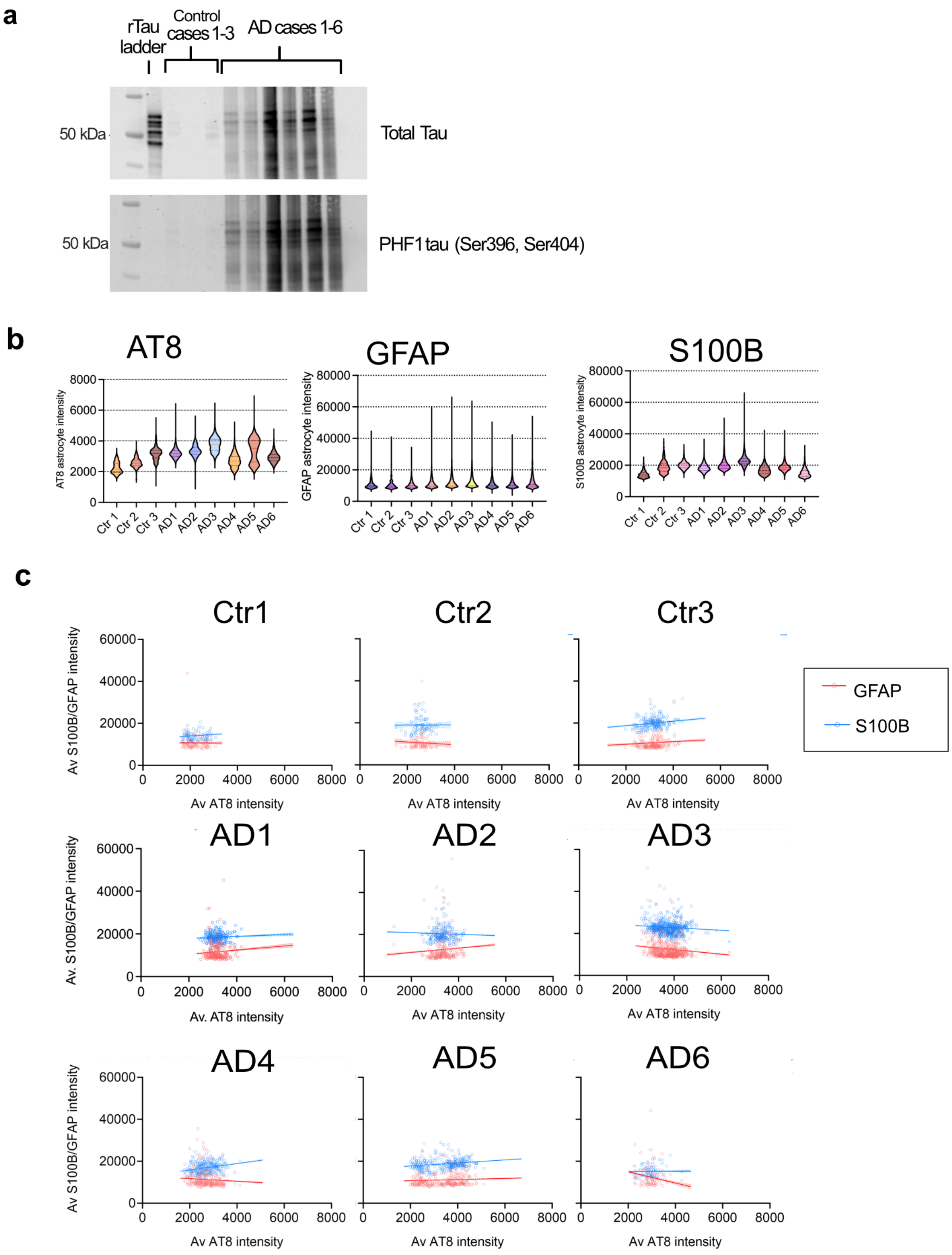


| | Ctr1 | Ctr2 | Ctr3 | AD1 | AD2 | AD3 | AD4 | AD5 | AD6 |
|---|---|---|---|---|---|---|---|---|---|
| **S100B** | | | | | | | | | |
| **r** | 0.1282 | 0.002558 | 0.191 | 0.05687 | -0.03765 | -0.09018 | 0.2348 | 0.2265 | 0.01587 |
| **$R^2$** | 0.01644 | 6.54E-06 | 0.03647 | 0.003234 | 0.001418 | 0.008132 | 0.05513 | 0.05128 | 0.000252 |
| **p** | 0.0002 | 0.9621 | <0.0001 | <0.0001 | <0.0001 | <0.0001 | <0.0001 | <0.0001 | 0.4377 |
| **GFAP** | | | | | | | | | |
| **r** | -0.00702 | -0.0603 | 0.09121 | 0.07605 | 0.06503 | -0.1111 | -0.07827 | 0.06584 | -0.1776 |
| **$R^2$** | 4.93E-05 | 0.003636 | 0.008319 | 0.005783 | 0.004229 | 0.01234 | 0.006126 | 0.004335 | 0.03153 |
| **p** | 0.837 | 0.2626 | <0.0001 | <0.0001 | <0.0001 | <0.0001 | <0.0001 | <0.0001 | <0.0001 |

### Supplementary Fig 2

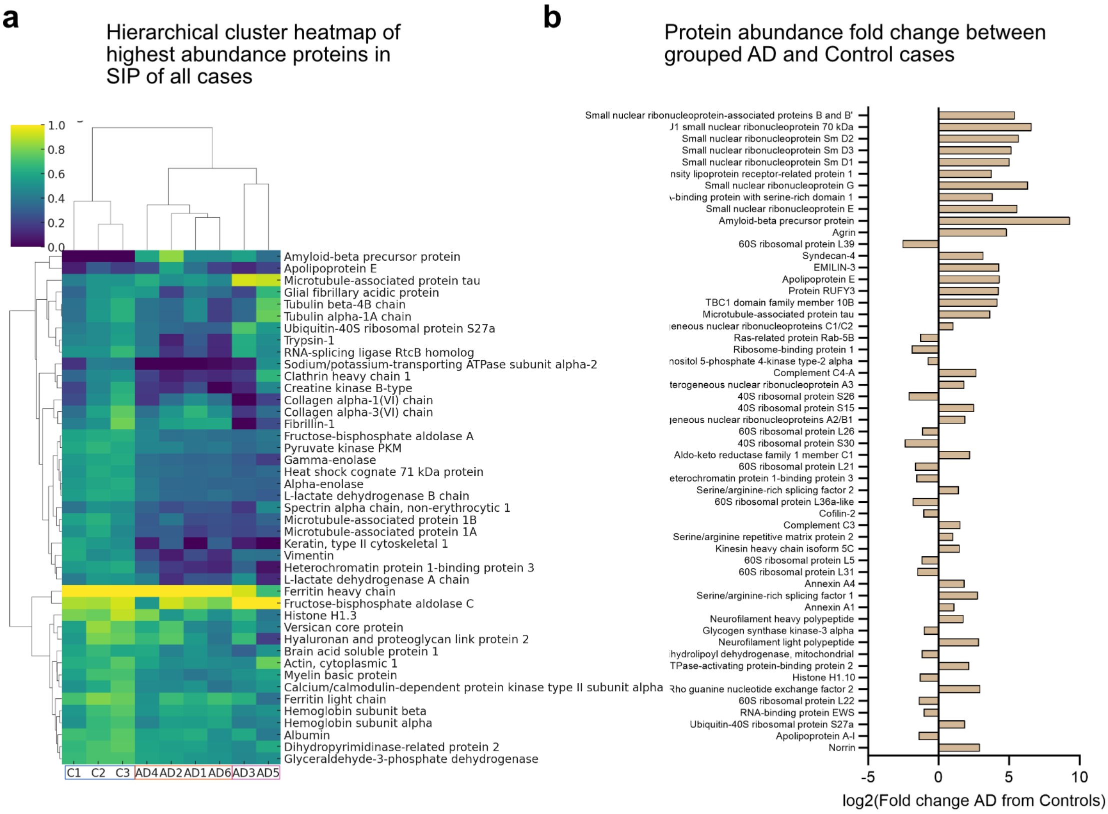

### Supplementary Fig 3

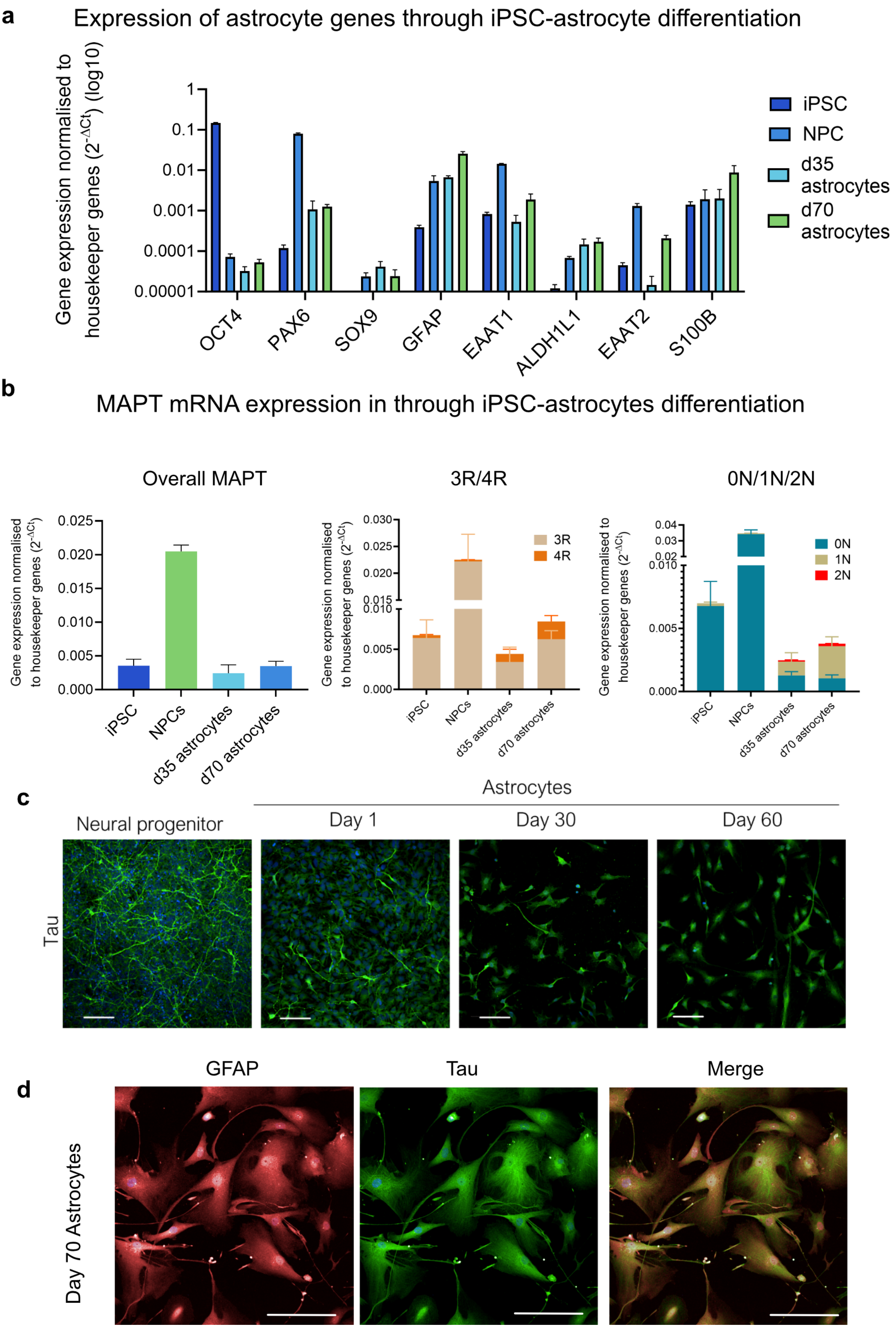

### Supplementary Fig 5

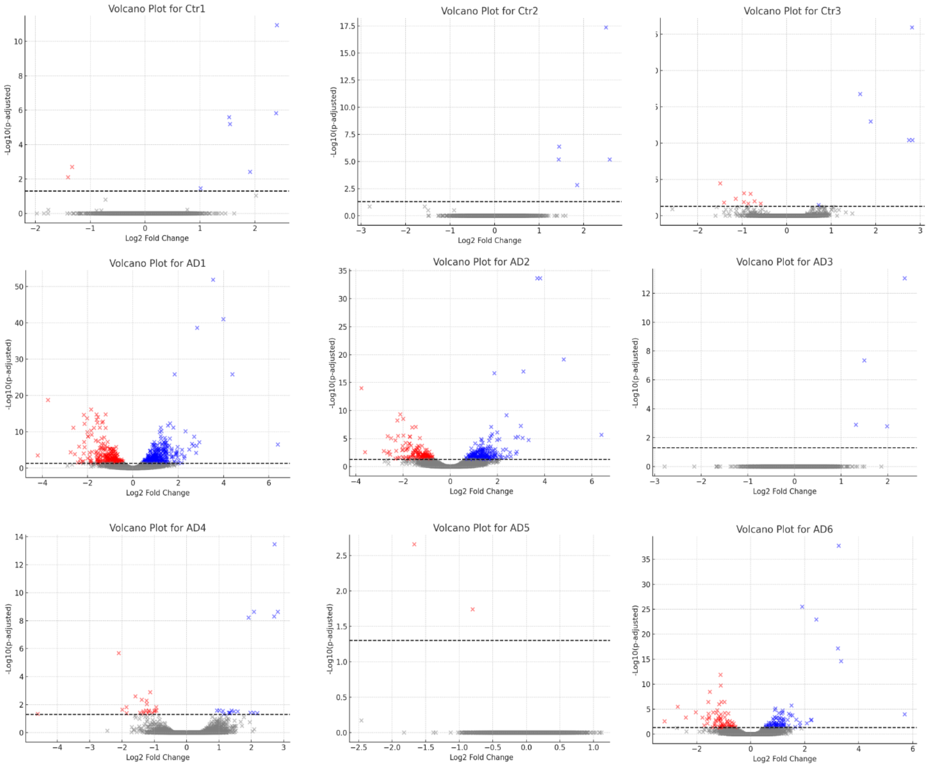

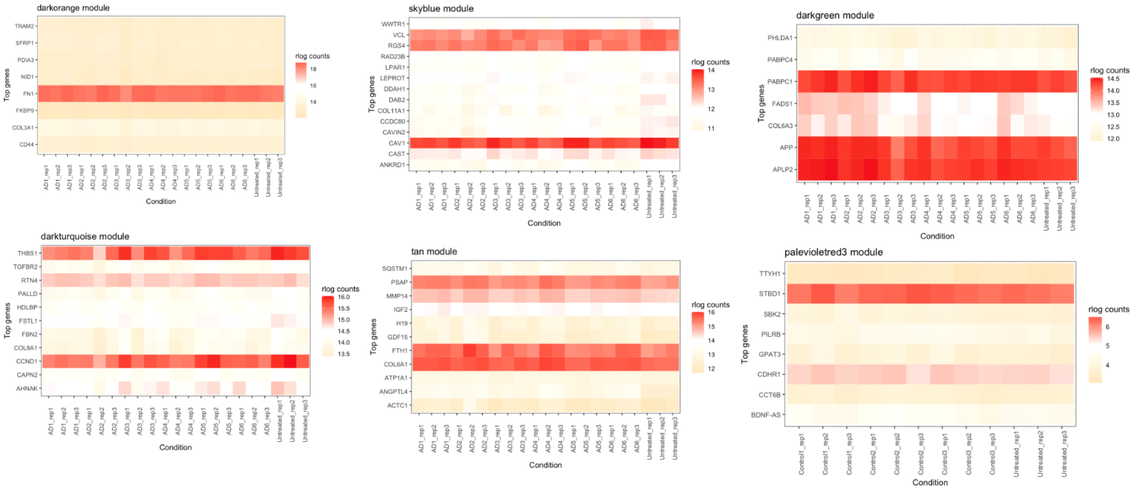
