## Supplementary Fig 4 for "Properties of Alzheimer’s disease brain-derived tau aggregates define tau processing by human astrocytes"

**a**

### Astrocyte AT8 vs GFAP/S100B intensity

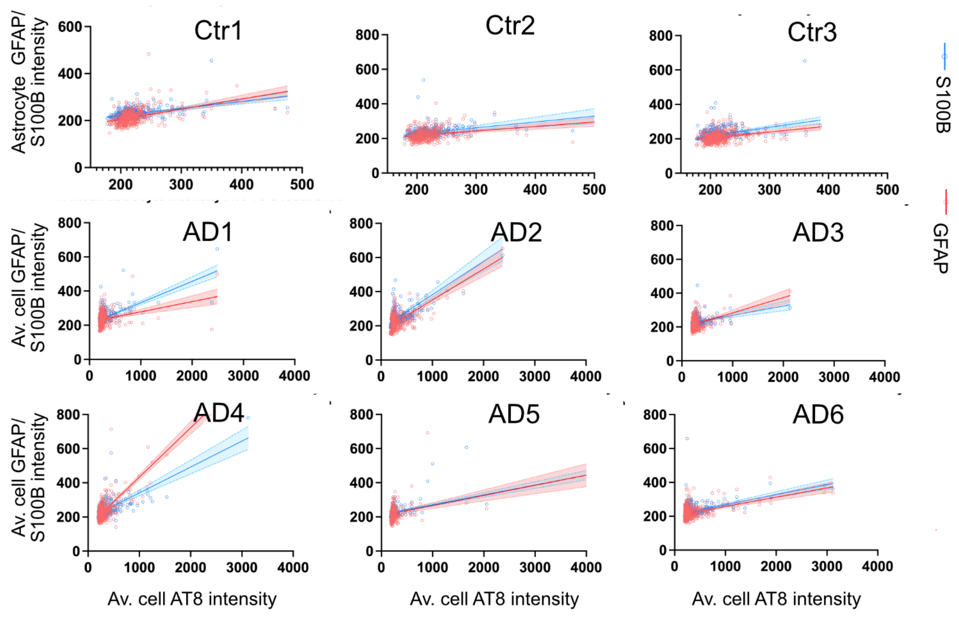


| | Ctr1 | | Ctr2 | | Ctr3 | | AD1 | | AD2 | | AD3 | | AD4 | | AD5 | | AD6 | |
|---|---|---|---|---|---|---|---|---|---|---|---|---|---|---|---|---|---|---|
| | S100B | GFAP | S100B | GFAP | S100B | GFAP | S100B | GFAP | S100B | GFAP | S100B | GFAP | S100B | GFAP | S100B | GFAP | S100B | GFAP |
| r | 0.4131 | 0.4377 | 0.2824 | 0.2179 | 0.275 | 0.3762 | 0.282 | 0.6174 | 0.5716 | 0.4761 | 0.3671 | 0.3207 | 0.7918 | 0.5243 | 0.3401 | 0.6626 | 0.4105 | 0.4616 |
| $R^2$ | 0.1707 | 0.1916 | 0.07977 | 0.04748 | 0.07561 | 0.1415 | 0.07953 | 0.3812 | 0.3267 | 0.2267 | 0.1347 | 0.1028 | 0.6269 | 0.2749 | 0.1157 | 0.4391 | 0.1685 | 0.2131 |
| *p* | <0.0001 | <0.0001 | <0.0001 | <0.0001 | <0.0001 | <0.0001 | <0.0001 | <0.0001 | <0.0001 | <0.0001 | <0.0001 | <0.0001 | <0.0001 | <0.0001 | <0.0001 | <0.0001 | <0.0001 | <0.0001 |

**b**

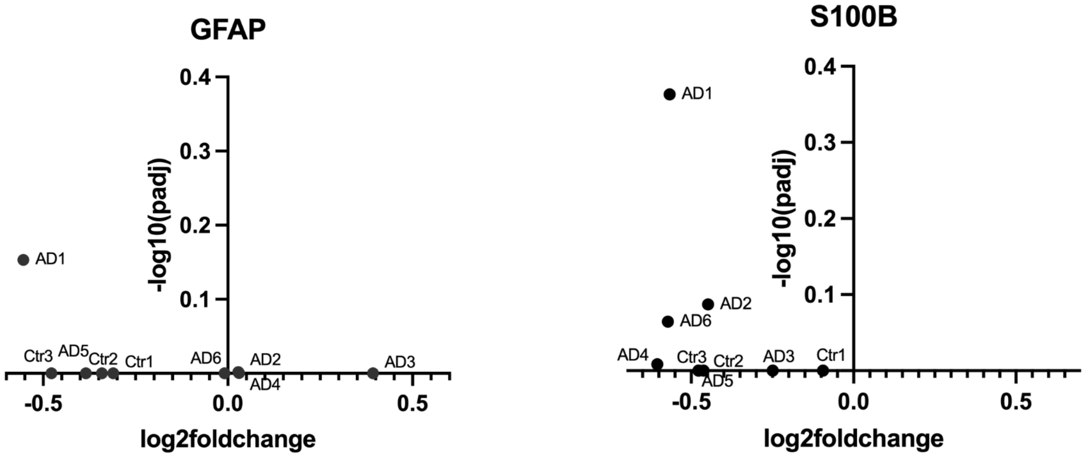


**c**

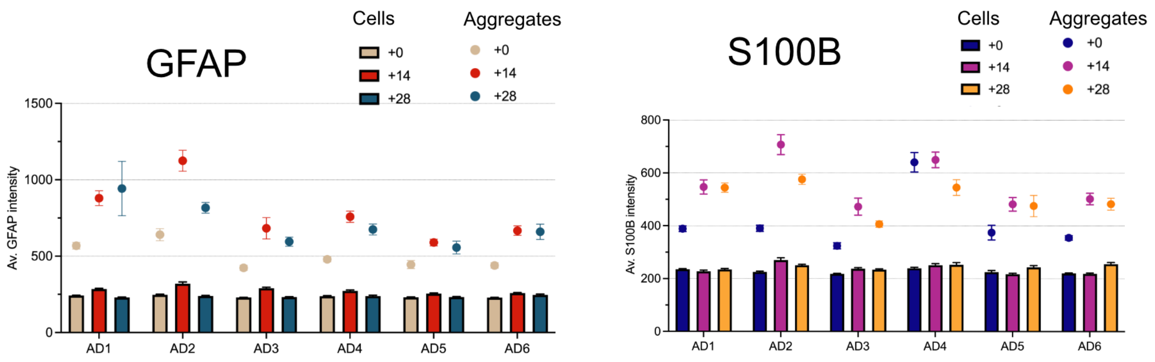
